# Disruption of the *Aspergillus fumigatus* RNA interference machinery alters the conidial transcriptome

**DOI:** 10.1101/2022.07.28.501871

**Authors:** Abdulrahman A. Kelani, Alexander Bruch, Flora Rivieccio, Corissa Visser, Thomas Krüger, Danielle Weaver, Xiaoqing Pan, Sascha Schäuble, Gianni Panagiotou, Olaf Kniemeyer, Michael J. Bromley, Paul Bowyer, Amelia E. Barber, Axel A. Brakhage, Matthew G. Blango

## Abstract

The RNA interference (RNAi) pathway has evolved numerous functionalities in eukaryotes, with many on display in Kingdom Fungi. RNAi can regulate gene expression, facilitate drug resistance, or even be altogether lost to improve growth potential in some fungal pathogens. In the WHO fungal priority pathogen, *Aspergillus fumigatus*, the RNAi system is known to be intact and functional. To extend our limited understanding of *A. fumigatus* RNAi, we first investigated the genetic variation in RNAi-associated genes in a collection of 217 environmental and 83 clinical genomes, where we found that RNAi components are conserved even in clinical strains. Using endogenously expressed inverted-repeat transgenes complementary to a conditionally essential gene (*pabA*) or a nonessential gene (*pksP*), we determined that a subset of the RNAi componentry is active in inverted-repeat transgene silencing in conidia and mycelium. Analysis of mRNA-seq data from RNAi double-knockout strains linked the *A. fumigatus* dicer-like enzymes (DclA/B) and RNA-dependent RNA polymerases (RrpA/B) to regulation of conidial ribosome biogenesis genes; however, surprisingly few endogenous small RNAs were identified in conidia that could explain this broad change. Although RNAi was not clearly linked to growth or stress response defects in the RNAi knockouts, serial passaging of RNAi knockout strains for six generations resulted in lineages with diminished spore production over time, indicating that loss of RNAi can exert a fitness cost on the fungus. Cumulatively, *A. fumigatus* RNAi appears to play an active role in defense against double-stranded RNA species alongside a previously unappreciated housekeeping function in regulation of conidial ribosomal biogenesis genes.

## INTRODUCTION

RNA interference (RNAi) is a conserved molecular mechanism that was likely intact in the last common ancestor of modern eukaryotes (Shabalina and Koonin 2008; Torri et al. 2022). RNAi uses small RNA (sRNA) molecules to suppress gene expression either transcriptionally by heterochromatin recruitment or post-transcriptionally through sequence-specific messenger RNA (mRNA) degradation or translational interference (Moazed 2009; Dang et al. 2011; Torres-Martinez and Ruiz-Vazquez 2017). The first reported example of RNAi in fungi came from the model fungus *Neurospora crassa* as a process termed quelling, where introduction of repetitive transgenes induced post-transcriptional gene silencing (Romano and Macino 1992). Since then, several RNAi and small RNA biogenesis pathways have been characterized in fungi that are largely classified as Dicer-dependent (canonical) RNAi or Dicer-independent (non-canonical) RNAi (Ketting 2011; Ruiz-Vazquez et al. 2015; Torres-Martinez and Ruiz-Vazquez 2017).

Dicer is a type IV RNase III enzyme responsible for dsRNA cleavage and biogenesis of a variety of sRNA classes in different organisms (Nicolas et al. 2015; Torres-Martinez and Ruiz-Vazquez 2017). In fungi, the orthologous RNase III domain-containing enzymes are typically referred to as dicer-like (DCL) proteins due to subtle differences in domain structure. The generated sRNAs are then loaded into an Argonaute protein, the main component of the RNA-induced silencing complex that binds and processes sRNAs to guide selective destruction, translational repression, or transcriptional suppression of target mRNAs (Agrawal et al. 2003; Torres-Martinez and Ruiz-Vazquez 2017). In some instances, RNA-dependent RNA polymerases are required for RNAi pathways to amplify the silencing activity by generating additional dsRNA substrates using the single-stranded sRNAs as primers (Torres-Martinez and Ruiz-Vazquez 2017). While the core RNAi proteins are mostly conserved across eukaryotes, some gene losses and duplications have occurred, which further diversified the RNAi pathways and their functional specialization (Hutvagner and Simard 2008; Mukherjee et al. 2013; Torres-Martinez and Ruiz-Vazquez 2017). Loss of RNAi was observed to fuel hypervirulence in *Cryptococcus* due to higher transposon activity (Gusa et al. 2020), and killer virus-infected *Saccharomyces cerevisiae* strains are able to kill and therefore outcompete non-infected strains in the absence of RNAi (Bostian et al. 1980). The RNAi machinery have also been implicated in other functions, ranging from ribosome biogenesis to maintenance of cellular quiescence (Bernstein et al. 2012; Roche et al. 2016), further highlighting the importance of RNAi across the fungal kingdom.

In this study, we improve our understanding of RNAi in the human fungal pathogen, *Aspergillus fumigatus. A. fumigatus* is a ubiquitous, filamentous saprophyte and among the leading fungal killers (Latge 1999; Beauvais et al. 2013). It encodes a functional RNAi system consisting of two dicer-like proteins (DclA and DclB), two argonautes (PpdA and PpdB), and two RNA-dependent RNA Polymerases (RrpA and RrpB), together with several other, annotated peripheral factors **(Fig. 1A)** (Mouyna et al. 2004; Hammond and Keller 2005; Hammond et al. 2008b). Here, we show that the *A. fumigatus* RNAi system is conserved among both clinical strains and environmental isolates. We then delineate the RNAi components necessary for inverted-repeat transgene (IRT) silencing and link the RNAi machinery to expression of conidial ribosome biogenesis genes and long-term fitness. Finally, we provide a brief characterization of the endogenous small RNA pool of *A. fumigatus*, which appears to be almost entirely separable from the observed differential expression and protein abundance in the knockout strains.

**FIG 1.**
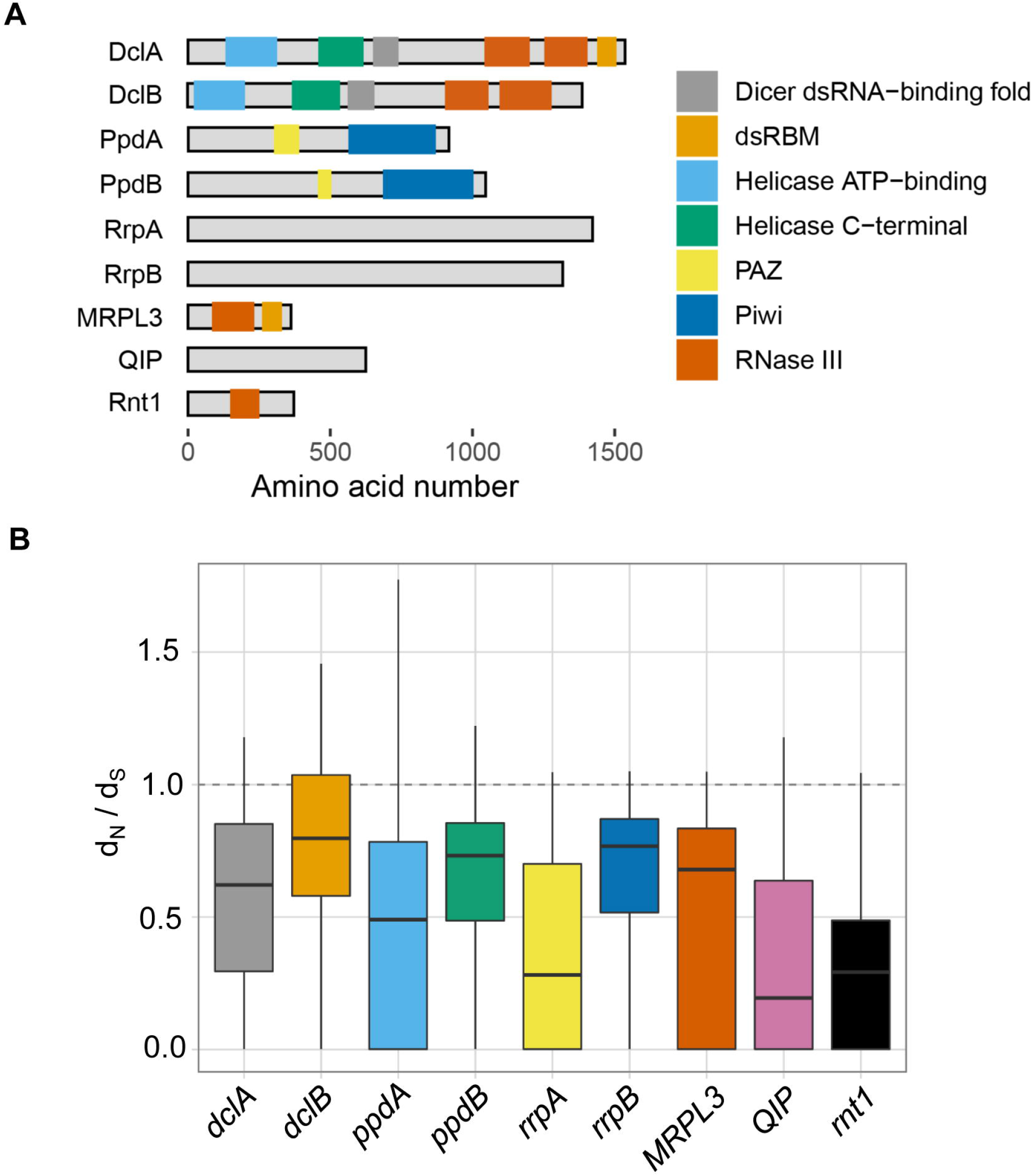
RNAi is conserved across 300 sequenced *A. fumigatus* strains. **(A)** Structural domains of *A. fumigatus* RNAi proteins were analyzed using UniProt (www.uniprot.org/). DclA and DclB both contain a DExH-helicase followed by a helicase-C, a Dicer-dsRNA binding fold that mediates binding with other proteins (Qin et al. 2010), a P-element-induced wimpy testis (PIWI)–Argonaute–Zwille (PAZ) domain for binding the *3’* end of the dsRNA, and two RNase III domains necessary for cleavage of bound dsRNA (Schauer et al. 2002; Mukherjee et al. 2013; Song and Rossi 2017). Both argonaute orthologs (PpdA/B) have the characteristic PAZ domain and a PIWI domain present in all eukaryotic argonautes (Jinek and Doudna 2009; Ender and Meister 2010). (B) Calculation of the ratio of nonsynonymous to synonymous base changes (d_N_/d_S_) for each of the core RNAi components and peripheral factors using 300 genomes from (Barber et al. 2021). Ratios under 1 indicate that each gene is under neutral-to-purifying selection.

## RESULTS

### RNAi pathway genes are conserved in environmental and clinical isolates of *A. fumigatus*

The RNAi pathway has diversified extensively since its emergence around the time of the last eukaryotic common ancestor (Torri et al. 2022). Functions range from gene regulation to genome defense, but in some specific instances the machinery has been lost to provide a growth advantage (Billmyre et al. 2013). To determine whether the RNAi machinery is stably maintained in *A. fumigatus, we* explored the core RNAi components for genetic variation across 300 *A. fumigatus* genomes collected from diverse clinical and environmental niches (Barber et al. 2020; Barber et al. 2021). All the RNAi pathway genes were present as a single copy in the genomes, indicating a high degree of evolutionary conservation. While many strains contained genetic variation, such as missense substitutions or alterations to the 3’UTR **(Fig. S1** and **Dataset S1),** high impact variants like frameshifts or premature stops were rare and only observed in 1-6% of samples (n= 4 for *dclA;* n = 1 for *rrpA;* n = 19 for *rrpB*). Of note, the *rrpA* and *rrpB* genes are thought to be dispensable for silencing in the related organism *Aspergillus nidulans* (Hammond and Keller 2005), whereas orthologs of the *dclA* gene contribute to the process of meiotic silencing of unpaired DNA during the sexual cycle in other organisms (Alexander et al. 2008). Calculation of the ratio of nonsynonymous to synonymous base changes (d_N_/d_S_) indicated that all members of the pathway are under purifying selection, often with one paralogous copy of the gene under a stronger degree of selection **(Fig. 1B).** Overall, we find a strong degree of genomic conservation among the *A. fumigatus* RNAi system, which together with the evidence of purifying selection suggests that the pathway likely plays a conserved, beneficial function across *A. fumigatus* strains and that loss does not strongly facilitate virulence in humans as has been hypothesized for other fungal pathogens (D’Souza et al. 2011; Billmyre et al. 2013).

### Engineered loss of the RNAi machinery provides no growth advantage in stress or drug challenge

Genomic analysis suggested that the RNAi machinery is not frequently lost under clinical or environmental conditions, but we wanted to explicitly test the hypothesis that RNAi is dispensable for stress response and drug resistance. To do this, we created a series of knockouts in the core RNAi machinery, including single knockouts of *dclA, dclB, ppdA, ppdB, rrpA*, and *rrpB* **(Fig. S2).** In addition, we made a series of pairwise knockouts in *dclA/B, ppdA/B*, and *rrpA/B*. None of the knockouts showed growth defects on minimal or rich media **(Fig. 2A).** Next, we examined the function of RNAi components in response to various stress conditions, including the cell wall stressors congo red, caffeine, and caspofungin; cell membrane stress agent sodium dodecyl sulfate (SDS); and oxidative stressor H_2_O_2_ **(Fig. S3).** We also tested for sensitivity to azole drugs like voriconazole and tebuconazole, as well as thiabendazole (TBZ), which is a microtubule inhibitor previously used to link RNAi components to chromosome segregation in fission yeast (Hall et al. 2003). In each case, the RNAi knockout strains showed no obvious phenotypic difference in comparison to the wild type.

**FIG 2.**
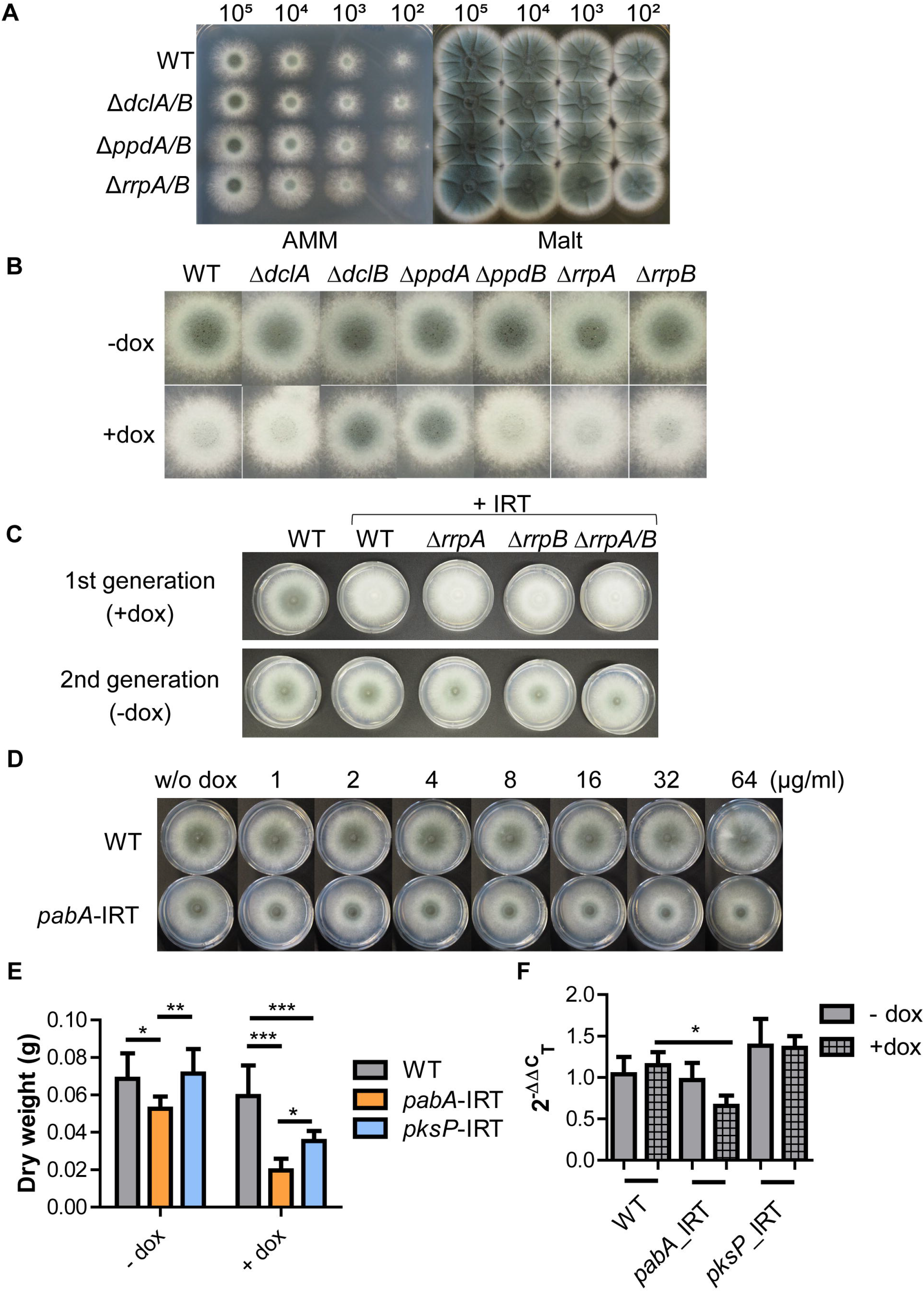
DclB and PpdA are required for IRT-mediated silencing in *A. fumigatus*. **(A)** Droplet growth assays for wild-type (WT), Δ*dclA/B*, Δ*ppdA/B*, and Δ*rrpA/B* strains grown on AMM or Malt agar plates for 48 h. Image shown is representative of three independent experiments. (B) Wild-type and RNAi knockout strains transformed with IRT constructs were spot inoculated on AMM with and without inducer (10 μg/ml doxycycline, **dox)** for 2 days. Image shown is representative of three independent experiments. (C) Wild-type and IRT-transformed strains were spot inoculated on AMM agar plates without inducer (10 μg/ml doxycycline) for 2 days. Spores were collected and the experiment was repeated without IRT induction. (D) The wild-type (CEA17Δ*akuB^KU80^*) strain was transformed with a plasmid (pSilent-pαbA) expressing a dox-inducible IRT targeting *pabA*, a para-aminobenzoic acid synthetase-encoding gene. Wild-type and *pabA*-IRT expressing strains, as well as a control *pksP*-IRT expressing strain were grown with and without dox induction for 24 h. Mycelia were collected and dried at 60°C for 3 days before measuring weight. (E) Wild-type and *pabA*-IRT expressing strain were spot inoculated on AMM agar plates supplemented with different concentrations of dox. Images were taken after 2 days of incubation at 37°C. Plot displays data from 4 biological replicates, with each replicate having 3 technical repeats. Significance was determined by two-way ANOVA with Bonferroni-corrected post hoc t-test comparing all columns. (F) Validation of silencing of *pabA* gene after induction of *pabA*-IRT using qRT-PCR. Plot shows standard error of the mean of three biological replicates performed in technical triplicates using the 2^-ΔΔC^_T_ method (Livak and Schmittgen 2001). Significance was determined by one-way ANOVA followed by Bonferroni corrected post hoc t-test comparing all columns and assuming Gaussian distributions.

### DclB and PpdA are necessary for inverted-repeat transgene silencing

The lack of phenotypic differences upon deletion of RNAi components led us to question the functionality and redundancy of the individual RNAi proteins in *A. fumigatus* and determine which core RNAi factors of *A. fumigatus* are required for silencing. It is known that exogenous delivery of RNA directly into the fungal cytoplasm is challenging due to a sequestration of negatively charged RNA by the cationic cell wall (Jochl et al. 2009). We first confirmed this finding in our own system using confocal microscopy of an exogenously delivered, 500-bp, fluorescein-labelled *in vitro*-transcribed dsRNA after incubation for 16 h at 37°C **(Fig. S4A).** The dsRNA clearly accumulated in the cell wall of the fungus co-stained with calcofluor white as expected.

Next, we turned to heterologous expression of two inverted-repeat transgenes from the *A. fumigatus* chromosome to target a nonessential gene and a conditionally essential gene **(Fig. S4B).** We reengineered a previously described system (Mouyna et al. 2004) using the pSilent-1 gene silencing vector (Nakayashiki et al. 2005) to produce a wild-type strain harboring a 500-bp, Tet-inducible *pksP or pabA* inverted-repeat transgene on the *A. fumigatus* chromosome at the *pyrG* locus. The nonessential *pksP* gene is required for dihydroxynaphthalene (DHN)-melanin biosynthesis, and silencing of *pksP*, results in an easy phenotypic readout via the production of pigmentless, white conidia (Langfelder et al. 1998). The second inverted-repeat transgene was complementary to the *A. fumigatus pabA* gene, which encodes a para-aminobenzoic acid synthetase (PabA). PabA catalyzes the biosynthesis of para-aminobenzoic acid (PABA), a precursor for folate biosynthesis in the fungus. Previous work has shown that deletion of *pabA* in *A. fumigatus* and *A. nidulans* results in loss of virulence and severe growth reduction, respectively, lending credence to *pabA* as a potential drug target (Brown et al. 2000). In fact, *A. fumigatus pabA* is conditionally essential, and only grows with appropriate supplementation (e.g. folate or p-aminobenzoic acid).

We first tested the *pksP* inverted-repeat transgene in the wild-type strain, where we observed white conidia after induction with doxycycline on AMM plates, indicative of successful post-transcriptional silencing of the *pksP* gene **(Fig. 2B).** Ultimately, we created two versions of these plasmids with different resistance cassettes so that we could also incorporate the construct onto the chromosome in each of our single and double knockouts. While similar silencing was observed in most transformed knockout strains, Δ*dclB* and Δ*ppdA* remained able to produce melanized conidia, suggesting that both DclB and PpdA are essential for inverted-repeat transgene silencing and cannot be rescued by their respective paralogs, DclA and PpdB.

As mentioned, the RdRPs contribute to the RNAi pathway in some organisms by transcribing additional dsRNA substrates and amplifying the silencing response (Dang et al. 2011). To eliminate the possibility that the overexpression of our inverted-repeat transgene RNA negated the necessity of the amplification role of RdRPs in *A. fumigatus, we* also passaged the induced RdRP strains onto fresh plates lacking inducer. This experiment allowed us to determine whether the silencing persisted after the induction of the inverted-repeat transgene was removed. In the wild-type, Δ*rrpA*, and Δ*rrpB* strains silencing was quickly lost **(Fig. 2C),** indicating that these proteins likely do not contribute to amplification of silencing in *A. fumigatus* and in agreement with results observed previously in *A. nidulans* (Hammond and Keller 2005). A curious observation is that the loss of silencing occurs to a lower extent than that of wild type, perhaps suggesting an additional layer of complexity surrounding the RdRP proteins in *A. fumigatus*.

Using the in vitro-transcribed dsRNA with complementarity to the conditionally essential gene *pabA, we* observed no cessation of growth in our imaging experiments **(Fig. S4A),** consistent with numerous previous studies showing that only minimal silencing can be achieved by the addition of exogenous RNA to *Aspergillus* species (Khatri and Rajam 2007; Abdel-Hadi et al. 2011; Kalleda et al. 2013; Mousavi et al. 2015; Nami et al. 2017). To test if effective silencing of *pabA* could lead to growth repression, we again used a heterologously expressed dsRNA targeting the *pabA* gene from the chromosome of *A. fumigatus* as described above **(Fig. 2D).** We grew both the pαbA-IRT-transformed strain and the wild-type strain with and without doxycycline induction on AMM agar plates and liquid cultures. In the absence of doxycycline, the *pabA*-IRT led to minor but significant growth inhibition, likely due to leaky expression **(Fig. 2D-E).** Upon induction with doxycycline, growth of the *pabA*-IRT-transformed strain was reduced on both AMM agar plates and liquid cultures when compared with the wild-type strain **(Fig. 2D-E).** Growth was more clearly inhibited in liquid culture than on solid media, likely due to more homogenous delivery of the doxycycline inducer. As a control, we also included the *pksP*-IRT strain in our liquid culture experiments and found that this construct also had a growth-inhibiting effect, albeit significantly less than the *pabA*-IRT strain **(Fig. 2E).** As a control, we confirmed by qRT-PCR that the *pabA*-IRT strain specifically targets *pabA*, resulting in a modest 43% decrease in message **(Fig. 2F).** Collectively, these results indicated that *A. fumigatus* RNAi can be leveraged to silence both a nonessential gene (*pksP*) and a conditionally essential gene (*pabA*) in a DclB- and PpdA-dependent manner.

### mRNA-seq analysis suggests distinct contributions to conidial gene expression

The inverted-repeat transgene experiments suggested that only select components of the *A. fumigatus* RNAi machinery are required for transgene silencing. We were therefore interested to elucidate any additional roles of RNAi in *A. fumigatus* biology. To this end, we performed mRNA-seq on poly(A)-enriched RNA isolated from conidia and mycelium grown in liquid culture for 24 or 48 h **(Fig. S5),** as RNAi is known to serve a role in asexual reproduction in other filamentous ascomycetes (Gaffar et al. 2019). We determined the differentially expressed genes **(Dataset S2)** and found that each of the RNAi knockouts showed significant differences compared to wild type in the conidia samples but very few changes in expression in mycelium using this stringent cutoff **(Fig. 3A).** These findings were consistent with the lack of phenotypes observed in stress and growth assays. We performed Gene Ontology (GO)-slim analysis to improve our understanding of these changes in conidia and were surprised to see a clear link between the *dcl* and *rrp* genes with ribosome biogenesis and the nucleolus, where 85 genes associated with ribosome biogenesis were significantly downregulated in the Δ*dclA/B* strain and 24 genes were downregulated in the Δ*rrpA/B* strain compared to wild type **(Fig. 3B; Fig. S6; Dataset S3).** The argonaute proteins appeared to contribute to conidial viability differently, as we observed decreased expression of genes associated with transmembrane transport and the regulation of transcription, alongside upregulation of genes associated with mRNA and tRNA metabolic processes.

**FIG 3.**
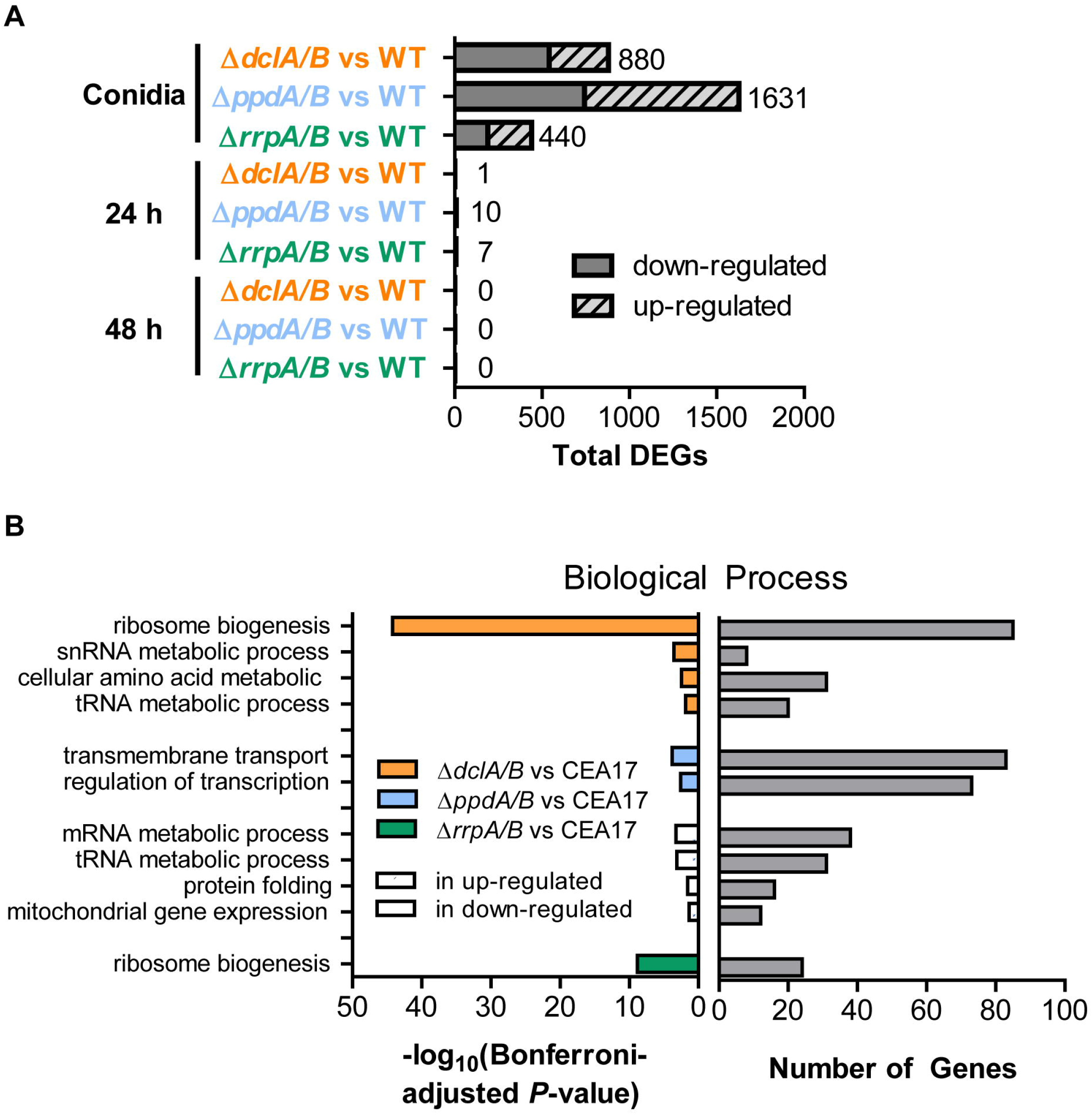
Deletion of RNAi components results in altered conidial transcriptome. **(A)** Genes that were identified as significantly differentially expressed in mRNA-seq. Differences were considered significant if q ≤ 0.05 and |log_2_ (MRN)| ≤ 1 by all four tools. The number next to the bar indicates the total number of differentially expressed genes (DEGs). (B) GO-Slim analysis (using FungiDB.org) for Biological Process generated using differentially expressed genes from conidia as input gene sets. Up- and down-regulated differentially expressed genes (left panel) as well as the number of differentially expressed genes (right panel) are indicated.

To expand on our mRNA-seq results, we performed LC-MS/MS-based proteomics on conidia and 48-hour-old mycelium for each of the double knockouts and the wild type. Among the most abundant proteins detected in conidia were several histones (HhfA, Htb1), a Cu/Zn superoxide dismutase (SodC), and elongation factor 1-alpha (AFUA_1G06390; **Dataset S4).** At 48 h, we observed a high abundance of alcohol dehydrogenase (AFUA_7G01010), the same Cu/Zn superoxide dismutase (SodC) as in conidia, and several ubiquitin isoforms (AFUA_1G04040 and AFUA_3G11260); results that are consistent with the metabolic changes associated with carbon and iron starvation (Oberegger et al. 2000) **(Dataset S5).** The dataset from conidia identified roughly 5,810 proteins, likely making this one of the most thorough descriptions of the proteome of *A. fumigatus* conidia to date. For the 48-h mycelium, we identified 2,380 proteins. Interestingly, from the top 200 most-abundant proteins from each dataset, we only observed 26 in common, reiterating the massive changes that are undergone during the *A. fumigatus* asexual life cycle.

We next determined the differentially abundant proteins between knockout and wild type using a two-tailed Welch’s t-test with Benjamini-Hochberg correction to control for multiple comparisons, which is known to be a stringent statistical test for proteomics analysis (Pascovici et al. 2016). The use of the Benjamini-Hochberg correction resulted in only three significant changes (adjusted P-value ≤ 0.05), all in the conidia condition **(Dataset S4).** Two proteins had lower abundance in the Δ*dclA/B* strain (AFUA_4G10100, AFUA_4G06250) and one protein (AFUA_7G08590) showed slightly increased abundance in the Δ*ppdA/B* strain compared to wild type **(Dataset S4).** The changes in AFUA_4G06250, a predicted ribosome biogenesis protein orthologous to Nop4, and AFUA_7G08590, a protein with a predicted BTB/POZ domain, were consistent with the change observed in our mRNA-seq analysis.

We also reperformed the statistical analysis using only a two-tailed Welch’s t-test, which revealed more widespread alterations to protein abundance in both conidia and 48-h-old mycelium **(Fig. 4, Fig. S7, Dataset S4-5).** In conidia, we observed 164 proteins with at least two-fold increase in abundance over wild-type in the Δ*dclA/B* strain and 113 proteins that significantly decreased in abundance. The Δ*ppdA/B* and Δ*rrpA/B* strains showed fewer significantly altered proteins, with 49 proteins increased and 49 decreased in abundance compared to wild type for the Δ*ppdA/B* strain and 37 proteins increased and 70 decreased for Δ*rrpA/B* **(Dataset S4).** Interestingly, we observed very little agreement between the mRNA-seq and proteomics analysis, implying that the conidial transcriptome differs substantially from the conidial proteome. In 48-h-old mycelium, we again observed significantly different protein abundances using a two-tailed Welch’s t-test. In particular, we found 35 proteins to be significantly more abundant (> 2-fold change) in the Δ*dclA/B* strain than wild type and 19 with decreased abundance; 61 proteins increased and 34 decreased in the Δ*ppdA/B* strain compared to wild type; and 86 proteins increased and 36 decreased in abundance in Δ*rrpA/B* **(Dataset S5).** Despite numerous cases of differential abundance, no clear pattern of regulation emerged from this analysis, potentially consistent with the idea that RNAi is pleotropic in its contribution to gene regulation. Collectively, these results suggest that the RNAi machinery in *A. fumigatus* influences fungal gene expression in multiple ways, with a particularly strong effect on genes associated with ribosome biogenesis in conidia.

**FIG 4.**
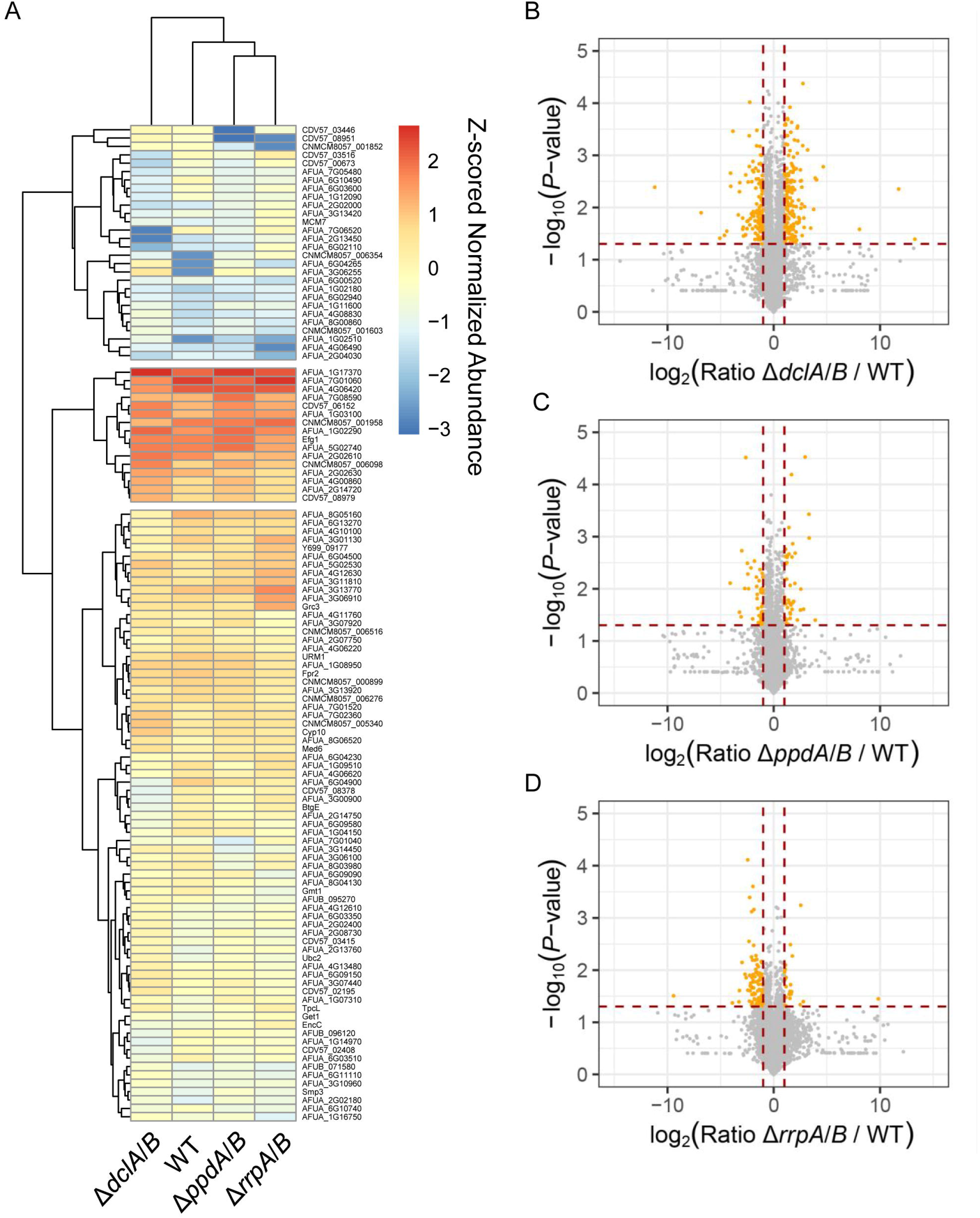
Conidia exhibit an altered proteome different from the transcriptome. (A) Clustered heatmap depicting Z-scored normalized average abundance data from conidia of double knockouts and wild type for cases where proteins exhibited at least a log_2_(FC)=2 and a P-value ≤ 0.05 in one of the conditions. Normalized abundances were determined using area under the curve from label-free quantification. (B-D) Volcano plots showing expression ratio for double knockout compared to wild type for (B) Δ*dclA/B*, **(C)** Δ*ppdA/B*, and (D) Δ*rrpA/B*. The y-axis shows the −log_10_ transformed *P*-value, where higher values are more significant. Orange circles show proteins that are at least log_2_(FC)=1 and with a P-value ≤ 0.05.

### RNAi genes exhibit growth stage-specific expression

The significant changes observed in conidia and mycelium led us to assess the expression of the different core and accessory RNAi components more closely in conidia versus mycelium for the wildtype and double-knockout strains using the data collected in the mRNA-seq experiment. The results clearly showed growth stage-specific expression for much of the RNAi machinery **(Fig. 5).** The *dclA* gene was more highly expressed in conidia than mycelium, whereas *dclB* increased with the age of the mycelium in a pattern similar to the mitochondrial ribosomal protein L3 (*MRPL3*) gene—known to encode an additional RNase III activity. The *rnt1* gene, which encodes a nuclear RNase III enzyme that could potentially compensate for the dicer-like RNase III activity in some instances, showed higher normalized expression in conidia than mycelium like *dclA*. The *ppdB* gene was expressed in all stages, whereas *ppdA* was essentially absent in conidia, consistent with a predicted role in post-transcriptional gene silencing that is likely unnecessary in relatively transcriptionally dormant conidia. Instead, we observed high expression of the exonuclease *QIP* in conidia, and not in mycelium, possibly suggesting QIP contributes to asexual as well as sexual reproduction as in other systems (Xiao et al. 2010).

**FIG 5.**
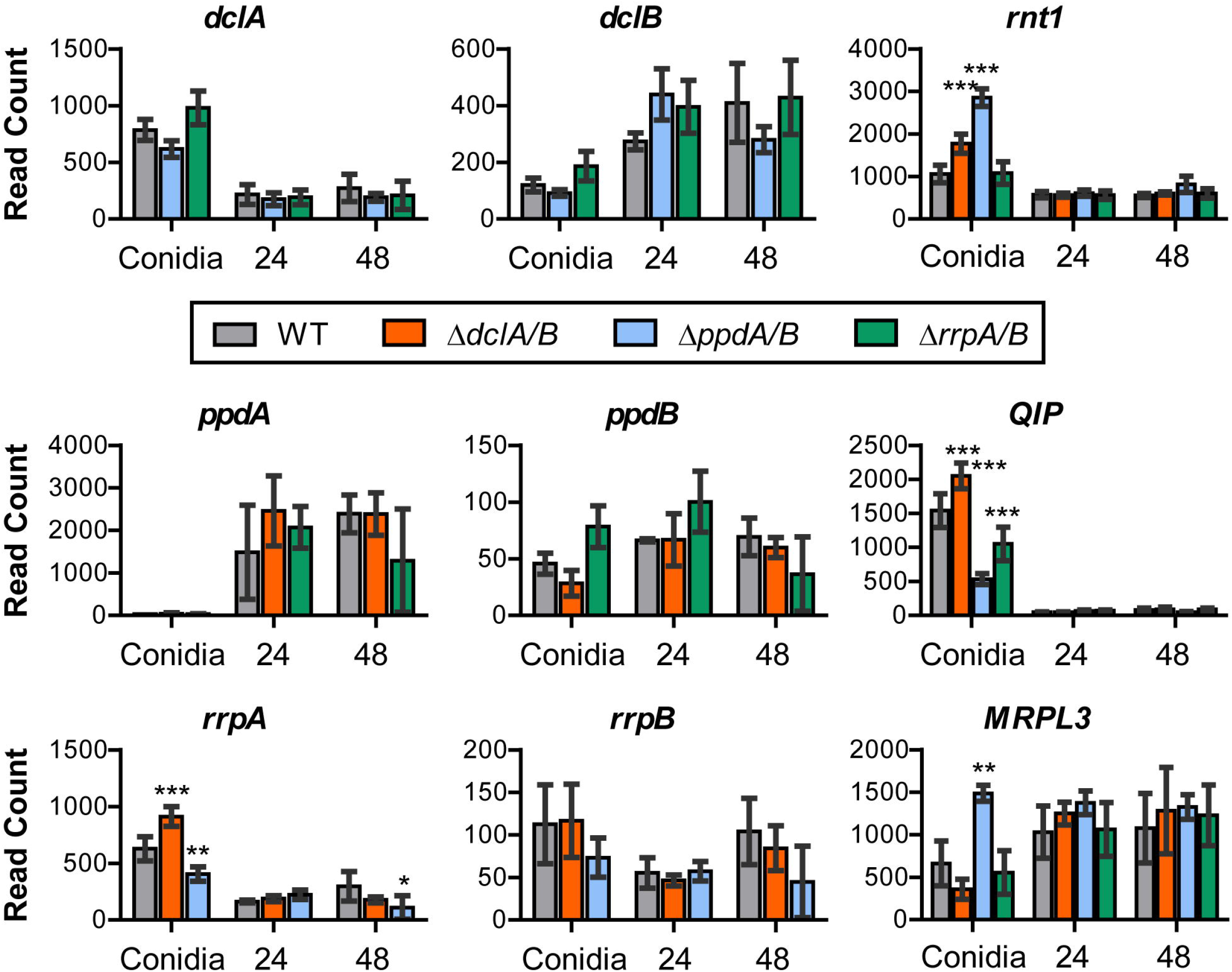
The RNAi genes are differentially expressed in conidia and mycelium. Median ratio normalization (MRN)-expression indicated as read count of individual genes in conidia, 24-h, and 48-h mycelium using the mRNA-seq experiment described in **Fig. 3.** Significance was determined by two-way ANOVA with Bonferroni-corrected post hoc t-test, resulting in some minor differences from the data analyzed in **Fig. 3**. *, P ≤ 0.05; **, P ≤ 0.01, ***, P ≤ 0.001.

Intriguingly, expression of many of the RNAi genes was significantly altered in the knockout strains as determined by two-way ANOVA with a Bonferroni posttest, with a few particularly notable examples. First, in the absence of the *dcl* or *ppd* genes, *rnt1* levels were significantly increased in conidia **(Fig. 5),** perhaps suggesting some level of compensation by this alternative RNase III enzyme. The expression of *QIP* and *rrpA* increased in the *dcl* double knockout in line with a known interaction between *QIP* and *rrpA*, whereas expression of both decreased in the *ppd* double knockout. Finally, *MRPL3* was greatly increased in conidia of the Δ*ppdA/B* strain, potentially connecting the argonaute proteins to regulation of translation in mitochondria and consistent with the observation of an increase in GO-slim genes associated with mitochondrial gene expression.

### RNAi contributes to fungal fitness over time

Next, to determine if the differential expression and protein abundance observed in the mRNA-seq and proteomics experiments might influence the long-term fitness of *A. fumigatus* RNAi knockouts, we performed serial passaging of the Δ*dclA/B, ΔppdA/B*, and Δ*rrpA/B* double knockouts and a wild-type control strain for 6 generations on AMM agar plates as separate lineages. Spores were spotted in the center of the plate and allowed to grow at 37°C for 7 days with minimal disturbance to provide numerous rounds of starvation and sporulation to recapitulate the asexual lifecycle of *A. fumigatus*. One sublineage of each of the RNAi knockout strains Δ*dclA/B* and Δ*ppdA/B* displayed subtle phenotypes that included white sectoring and altered production of conidia **(Fig. S8).** Previous reports have indicated that these white colonies typically result from mutation of the *pksP* gene, which encodes a polyketide synthase enzyme essential for the initial step in the biosynthesis of DHN melanin (Langfelder et al. 1998; Gibbons et al. 2022). Conidia collected from the white sectors in our experiment remained white upon transfer to new plates and were confirmed by PCR to hold the appropriate resistance cassette of the knockout. We then counted spore concentrations after a final passaging and observed a decrease in spore abundance from several lineages of the Δ*dclA/B* and Δ*ppdA/B* double knockouts **(Fig. 6),** suggesting that the Δ*dclA/B* and Δ*ppdA/B* knockouts can become less reproductive over time.

**FIG 6.**
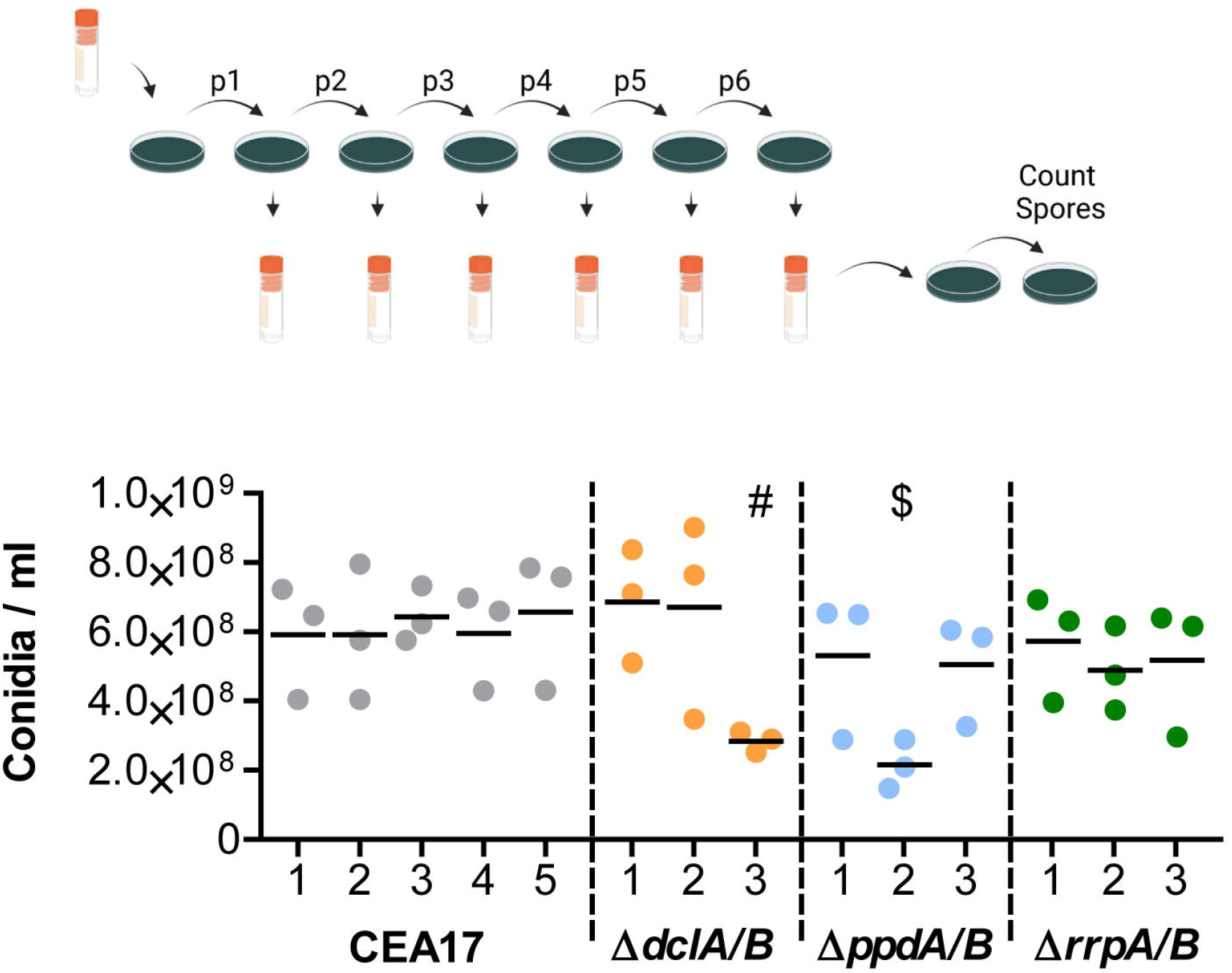
Deletion of RNAi components affects long-term fitness. Wild-type and double knockout strains were grown as >3 lineages and passaged for 6 generations on AMM agar plates at 37°C for 7 days (top). Model Created with Biorender.com. The evolved strains were then grown for an additional week and the spores were counted using a CASY counter as three biological replicates. The plot shows the values from each biological replicate of each lineage and the mean value of the biological replicates (black line). Significance was determined by one-way ANOVA with Bonferroni-corrected post hoc t-test comparing all columns. # indicates P-value ≤ 0.05 and $ indicates P-value ≤ 0.01 for all five comparisons with wild-type lineages.

### *A. fumigatus* produces a limited pool of high-abundance endogenous small RNAs

Finally, we assessed the endogenous pool of small RNAs generated in *A. fumigatus* CEA10 wild type to uncover potential small RNAs with complementarity to the observed differentially expressed genes and proteins. We performed small RNA sequencing using RNA again isolated from resting conidia and mycelium grown for 24 or 48 h in liquid AMM culture. The RNA was processed with RNA 5’ pyrophosphohydrolase (RppH) to remove any 5’ triphosphates before library preparation to facilitate small RNA discovery. The resulting library concentrations were quite low for resting conidia and mycelium grown for 24 h, so we did not normalize the library input before sequencing, and instead sequenced all the available libraries to maximize sRNA discovery. Not surprisingly, we found the largest number of reads from the 48-h mycelium samples where we also observed the largest amount of small RNA in in preparative analyses **(Table S1).** We predicted small RNAs using Manatee, a web-based algorithm for identification and quantification of sRNAs (Handzlik et al. 2020). 327 sRNAs were initially predicted by the program for the three timepoints (conidia, 24-h mycelium, 48-h mycelium; **Dataset S6).** 42 small RNA regions were identified in the conidia dataset, 77 in the 24-hour data, and 208 from the 48-hour data. Removal of duplicated regions found in multiple samples resulted in 297 unique identifications, with 14 found in two conditions and 8 found at all three timepoints **(Dataset S6).** After applying a cutoff to remove transcripts with fewer than 200 total reads, we were left with a total of 23 high-abundance sRNA candidates (**Table** 1). These small RNA loci range from 19 to 25 nucleotides in length, although the exact 5’ and *3’* nucleotides were not experimentally defined. The lack of abundant candidates is consistent with previous work on *A. fumigatus* endogenous small RNAs (Ozkan et al. 2017).

**Table 1.**
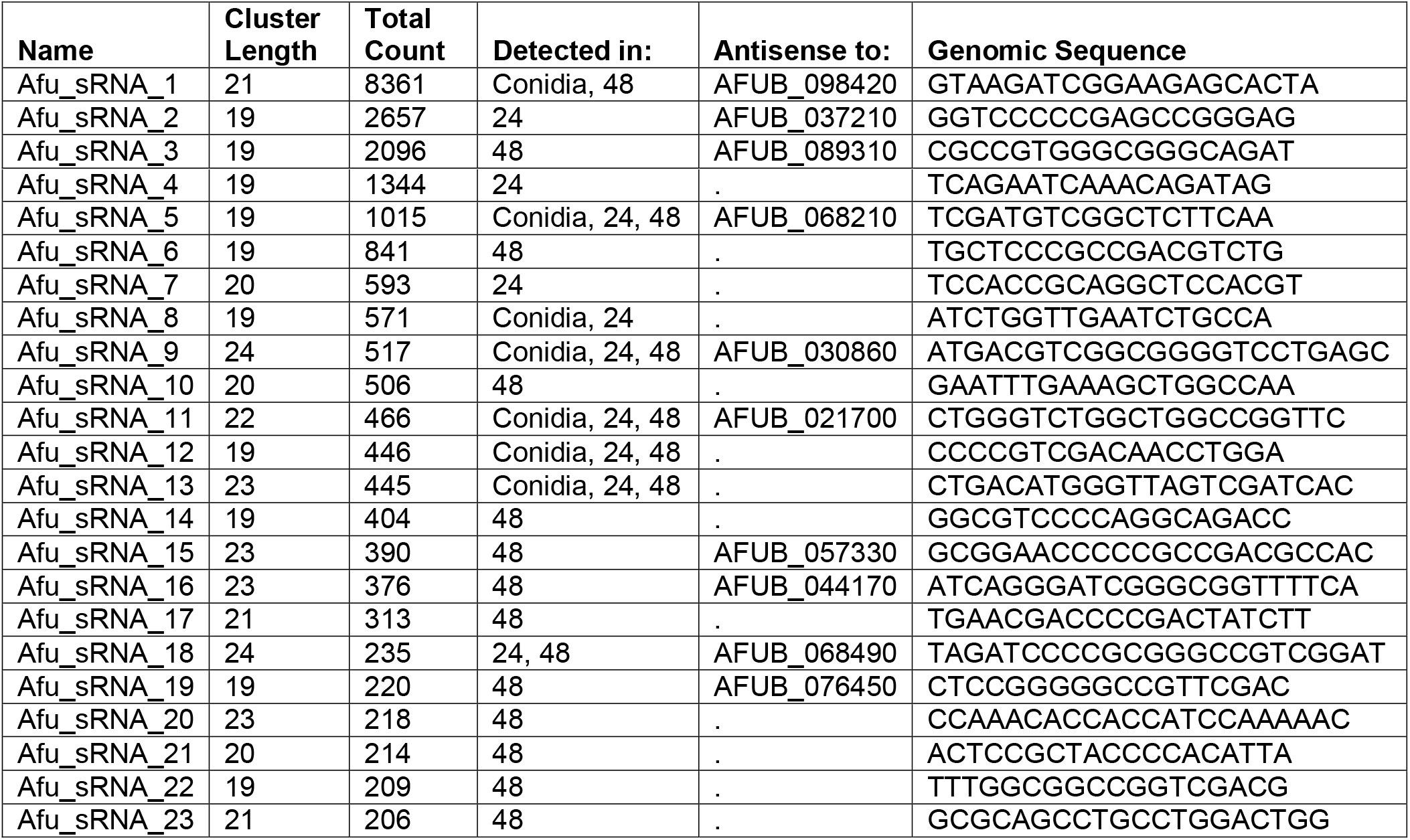
*A. fumigatus* Smalls RNAs identified by Manatee with over 200 reads.

The most abundant small RNAs in conidia were antisense to AFUB_098420 (AFUA_4G04590), an uncharacterized gene predicted to be associated with ubiquinone biosynthesis, and AFUB_068210 (AFUA_4G11180), a predicted AAA family ATPase. As expected, we observed a higher abundance of small RNAs at 48 h, but these small RNAs also had limited complementarity to proteins with altered abundance in 48-h-old mycelium or conidia samples. Certainly, a more extensive survey of the small RNAs produced by *A. fumigatus* using the different RNAi knockout strains generated here will be required to ultimately tease apart the complex regulation across the asexual lifecycle.

## DISCUSSION

The *A. fumigatus* RNAi pathway is functional in post-transcriptional gene silencing and involved in defense against mycoviruses (Mouyna et al. 2004; Ozkan et al. 2017). While work done in *A. nidulans* laid the groundwork for understanding further RNAi in *A. fumigatus*, distinct differences between the organisms, including truncations of the A. *nidulans* proteins (Hammond and Keller 2005; Hammond et al. 2008a; Hammond et al. 2008b), suggested that additional experimentation could be beneficial. We were able to show that the truncations of some core RNAi components in A. *nidulans* are not commonly repeated in *A. fumigatus*, as 300 sequenced genomes of *A. fumigatus* showed only rare truncation events (e.g. frameshifts) and relatively minor genetic variation. This relatively high level of conservation coupled with the observation that modest silencing can result in growth inhibition **(Fig. 2F)** suggests that we may even be able to exploit this machinery for therapeutic purposes if we can better facilitate RNA delivery in the future, as is being pursued with agricultural pests (Cai et al. 2019; Bruch et al. 2021).

Using a previously established inverted-repeat transgene approach (Mouyna et al. 2004), we created an inducible dsRNA against the *pksP* gene of *A. fumigatus*, which is required for production of DHN-melanin and the grey-green color of conidia. With this approach, we were able to determine that DclB and PpdA are required for inverted-repeat transgene silencing in *A. fumigatus*, whereas the other core components are dispensable for this form of RNA silencing. The finding that DclB and PpdA are essential for silencing matches previous work in *N. crassa*, where DCL-2 and QDE-2, orthologs of DclB and PpdA, were shown to be important for quelling (Dang et al. 2011). The result that *A. fumigatus ppdA* is necessary for silencing also matches results from *A. nidulans* with ortholog RsdA (Hammond and Keller 2005). Unlike *N. crassa*, DclA was not able to compensate in a Δ*dclB* deletion strain of *A. fumigatus* in our *pksP*-inverted-repeat transgene silencing experiments, suggesting discrete functions for DclA, perhaps during the poorly understood *A. fumigatus* sexual cycle. QDE-1 is important for silencing in *N. crassa*, but lost in A. *nidulans* (Hammond and Keller 2005), whereas the ortholog encoded by *rrpA* is conserved, but dispensable for IRT silencing, in *A. fumigatus* **(Fig. 2B-C).** This result appears to be consistent with the described loss of the activity in A. *nidulans*.

We showed that a 500-bp double-stranded RNA with complementarity to the *pabA* mRNA is capable of silencing *pabA* expression to limit fungal growth when expressed from the chromosome. Surprisingly, we also observed that the *pksP*-IRT led to growth inhibition when heterologously expressed. We suspect that the *pksP*-IRT sequence either results in production of small RNAs that have off-target effects or that the overproduction of a double-stranded RNA is mistaken for a mycoviral infection, resulting in altered fungal behavior. It is possible that overexpression of double-stranded RNA, which may also mimic a mycoviral infection, sequesters RNase III proteins from their normal functions in ribosome biogenesis and limits fungal growth. It is worth noting that in the literature, heterologous overexpression of a much less structured human mRNA, RBP7, in *A. fumigatus* did not detrimentally alter growth (Shopova et al. 2020), suggesting that the observed growth inhibition is not due solely to resource depletion, e.g. nucleotide pool depletion. A closer inspection of the small RNAs produced from the inverted-repeat transgene constructs will be required to fully understand this observation and detail the interplay between the different functionalities of RNAi in *A. fumigatus*.

The RNAi pathway in fungi has been linked to diverse regulation, including in sexual differentiation, development, stress response, and growth (Ruiz-Vazquez et al. 2015; Nicolas and Garre 2016; Torres-Martinez and Ruiz-Vazquez 2017). Despite all these important functions in other organisms, *A. fumigatus* appears to be relatively robust in tolerating deletion of the core RNAi machinery. Minimal functional differences were observed in a wide array of stress assays, although we did observe a decrease in fitness in our serial passaging experiment for some lineages of the Δ*dclA/B* and Δ*ppdA/B* strains, implying that loss of RNAi can have consequences. RNAi pathways also frequently play an important role in maintaining genome integrity, as has been described in distantly related fungi *Cryptococcus neoformans* (Janbon et al. 2010; Wang et al. 2012), *Mucor circinelloides* (Nicolas et al. 2010; Calo et al. 2017), *Fusarium graminearum* (Chen et al. 2015), *Magnaporthe oryzae* (Kadotani et al. 2003; Murata et al. 2007), *Schizosaccharomyces pombe* (Roche et al. 2016), and *Trichoderma atroviride* (Carreras-Villasenor et al. 2013). Investigation of fungal RNAi has proven a function in preserving the integrity of the host genome against invasive nucleic acids including transgenes, transposons, and invading viruses during both the vegetative and sexual cycles (Nuss 2011; Torres-Martinez and Ruiz-Vazquez 2017). Future studies into the RNAi components that contribute to genome protection are now possible using the tools we created here.

Additional work will be needed to determine the mechanism of regulation imparted by the dicer-like proteins in conidia, but some clear differences from previous reports should be noted. For example, *A. fumigatus* has a conserved, seemingly intact Rnt1 enzyme and the action of the dicer-like proteins seems unlikely to match that observed in *C. albicans*, where the only active RNase III enzyme performs the function of both dicer and Rnt1 in processing of rRNA (Bernstein et al. 2012). RNAi has also been linked to maintenance of quiescence in S. *pombe*, but again differences exist between the two organisms. *A. fumigatus* RNAi knockouts showed no defects in growth in our assays, and no change was observed when the RNAi knockouts were stressed with TBZ. The effects in conidia could also be due to regulation of heterochromatin formation as has been shown in other organisms (Martienssen and Moazed 2015), but at least preliminary studies in *A. nidulans* suggest that RNAi is uncoupled from regulation of heterochromatin in *Aspergillus* (Hammond and Keller 2005).

In the proteomics analyses, we observed only minor differences using a strict statistical analysis, whereas a more relaxed analysis led to disparate changes. As RNAi is known to play pleotropic roles in gene regulation, we can envisage that *A. fumigatus* may use RNAi to control gene expression in a variety of different ways. This pleotropic regulation is the limited pool of endogenous small RNAs in *A. fumigatus*. The majority of reads from small RNA-seq were derived from ribosomal RNAs and tRNAs, as was observed previously (Jochl et al. 2009), with only 23 high-abundance small RNA loci. These matched poorly to observed changes in protein abundance in the knockouts, potentially suggesting alternative biogenesis pathways or regulation in the conidiophore structure. A more in-depth analysis using the deletion strains produced here will be required to understand the pool of small RNAs in *A. fumigatus*.

In conclusion, we have shown that *A. fumigatus* RNAi appears to play an active role in post-transcriptional gene silencing that runs parallel to a previously unappreciated housekeeping function in regulation of conidial ribosomal biogenesis genes. The strains created and datasets collected here will likely be useful in further delineating the functions of this complex, evolutionarily diverse RNA regulatory pathway.

## MATERIALS AND METHODS

### Domain mapping and genomic analysis

Domain structures were created using UniProt annotations (Uniprot.org; Q4WSH7, Q4WY64, Q4WEW6, Q4WWR4, Q4WUR8, Q4WC92, Q4WXX3, Q4WA22, and Q4WVE3) and created with the drawProteins (Brennan 2018) package using R version 4.1.0 (2021-05-18).

The genomic data analyzed in this study was generated in (Barber et al. 2021) and is publicly available as raw reads and assembled genomes under PRJNA697844 and PRJNA595552. Isolates originated from environmental and clinical sources and displayed a global distribution. Presence/absence variation in the genomes was quantified using Orthofinder using peptide sequences from 300 genomes. Given that no cases of gene loss or amplification were detected among the genome assemblies, analysis of sequence variation was performed by aligning raw sequencing reads to the Af293 reference genome (FungiDB release 39). PCR duplicates were marked using MarkDuplicate from Picard v.2.18.25. Variant calling to detect SNVs and short indels was performed using HaplotypeCaller and high-quality variants identified using the following filtering criteria: SNP: QD < 2.0 || MQ < 40.0 || FS > 60.0 || MQ RankSum < −12.5 || ReadPosRankSum < −8.0; indel: QD < 2.0 || FS > 200.0 || ReadPosRankSum < −20.0. Variant function was predicted using SnpEff v.4.3.

### Strains and culture conditions

*A. fumigatus* strains were grown on *Aspergillus* minimal medium (AMM) agar plates (70 mM NaNO_3_, 11.2 mM KH_2_PO_4_, 7 mM KCl, 2 mM MgSO_4_, 1% (w/v) glucose and 1 μl/ml trace element solution (pH 6.5)). The trace element solution was composed of 1 g FeSO_4_ • 7 H_2_O, 8.8 g ZnSO_4_ • 7 H_2_O, 0.4 g CuSO_4_ • 5 H_2_O, 0.15 g MnSO_4_ • H_2_O, 0.1 g NaB_4_O_7_ • 10 H_2_O, 0.05 g (NH_4_)_6_Mo_7_O_24_ • 4 H_2_O, and ultra-filtrated water to 1000 ml. Spores were harvested after 5 days (or 7 days for the evolution experiment) with 10 ml of sterile distilled water using a T-shaped inoculation spreader and spore suspensions were filtered through a 30-μm cell strainer (MACS, Miltenyi Biotec GmbH, Germany) to exclude mycelium. *A. fumigatus* transformants were screened on AMM agar plates supplemented with 150 μg/ml of hygromycin, 0.1 μg/ml of pyrithiamine, or 160 μg/ml of phleomycin for selection. For *in vitro* transcription and confocal laser scanning microscopy experiments, *A. fumigatus* AfS150 was grown on malt agar (Sigma-Aldrich) plates supplemented with agar to a final concentration of 2% (wt/vol) for 5 days. *Escherichia coli* strains were grown at 37°C on LB broth or agar supplemented with 100 μg/ml carbenicillin when necessary.

### Genetic manipulation strategies

Knockout constructs were created using several strategies, namely Fusion PCR or the Gibson cloning method (9). To make constructs by Fusion PCR (10), an antibiotic selection gene and approximately 1 kb of DNA flanking regions (5’ and 3’) of the target gene were amplified. PCR fragments were purified using the GeneJET PCR Purification Kit (Thermo Fisher Scientific; Vilnius, Lithuania) and concentrations were measured by NanoDrop 2000 spectrophotometer. Next, equimolar amounts of the three fragments, two flanking regions, and selection gene, were combined using Fusion PCR with the 2x Phusion Flash High Fidelity Master Mix and 10 pmol of each forward and reverse primer for the 5’ flank and 3’ flank, respectively. Fusion PCR products were purified and validated by agarose gel electrophoresis. For Gibson assembly, a selection gene and about 1 kb flanking regions of target gene were amplified by PCR using primers that create overhangs **(Table S2).** In addition, pUC18 vector backbone was also amplified from plasmid “pUChph Volker”. DNA fragments were assembled and the resulting plasmid was used to transform chemically-competent *E. coli* following the NEBuilder HiFi DNA Assembly protocol (New England Biolabs; Ipswich, United States). Knockouts of *A. fumigatus* were generated by homologous recombination following the transformation of protoplasts (Weidner et al. 1998).

### Growth and viability assays

The growth capacity of different *A. fumigatus* strains in AMM media was investigated by determining the dry weight from shaking cultures. 1×10^7^ spores were inoculated in 50 ml AMM and incubated at 37°C and 200 rpm for 24 h. Mycelia were harvested using Miracloth (Merck KGaA; Darmstadt, Germany), washed with deionized water, and dried at 60°C for 3 days. The susceptibility of different *A. fumigatus* strains to stress conditions was investigated by spotting 5 μl of spore suspension with concentrations ranging from 10^5^-10^2^/ml on AMM agar plates supplemented with different stress-inducing agents for 2 days. All plates were incubated at 37°C.

### Inverted-repeat transgene silencing assays

To silence *A. fumigatus pksP*, sense and anti-sense *pksP* PCR product (500 bp each) were amplified with primer pairs oARK0031 & 0032 and oARK0033 & 0034 respectively. These PCR products were then cloned into the two multi-cloning sites in the p-Silent1 plasmid vector, which already has a hygromycin resistant cassette. The *trpC* promoter upstream the inverted-repeat region original present from the p-Silent 1 vector was replaced by a tetracycline inducible promoter (TetON). The resulting plasmid was then digested with restriction enzymes *SacI* and *FspI* to excise the IRT expressing region and hygromycin (hph) resistant cassette, which was then cloned into a pUC18 vector backbone flanked by 5’ and 3’ regions (approximately 1 kb) of a non-functional *pyrG* locus in the *CEA17*Δ*akuB^KU80^* (A1163, wild type) strain to generate pSilent-*pks*P1. Because some of the RNAi knockout strains already have a hygromycin resistant cassette, we constructed pSilent-*pks*P2 by replacing the selection marker with a phleomycin (ble) resistant gene. Both plasmids were linearized by *BtsI* restriction enzyme before transformation of strains **(Table S3).** To generate a para-aminobenzoic acid synthetase (*pabA*) silencing construct, *pabA*-IRT sequence plus a 3’ *trpC* terminator was synthesized. This synthesized DNA was then cloned using Gibson assembly into a PCR product amplified from pSilent-*pks*P2 with primer pair oARK0096 & oARK0038. The latter PCR product consist of the pUC18 backbone, *pyrG* flanks, phleomycin resistant cassette, and TetON promoter to drive the *pabA*-IRT expression. Plasmid was linearized by *PstI* restriction enzyme before transformation of *CEA17*Δ*akuB^KU80^* wild-type strain.

### Southern blot

Target gene deletion was confirmed by Southern blot, as described previously (Southern 2006). Digested genomic DNA samples were separated in a 1% (w/v) agarose gel. Separated DNA fragments on gel were transferred onto a nylon membrane (pore size: 0.45 μm, Carl Roth GmbH; Karlsruhe, Germany). The DNA was crosslinked to the membrane by UV irradiation followed by overnight hybridization with a DIG-11-dUTP-labelled DNA probe at 65°C. The membrane was washed twice at 65°C for 5 min (1x SSC, 0.1 SDS), followed by 2 washes with wash buffer II (0.5x SSC, 0.1 SDS) for 20 min at 65°C. The membrane was next rinsed with maleic acid wash buffer, incubated with blocking solution for 30 min to prevent unspecific antibody binding, followed by another incubation with 20 ml blocking solution plus 0.5 μl Anti-Digoxigenin-AP, Fab fragments (Roche, Mannheim, Germany) for 30 min. The unbound antibodies were removed by washing twice with 50 ml maleic acid wash buffer for 15 min. Finally, the membrane was equilibrated with detection buffer, developed with several drops of the chemiluminescent substrate CDP-Star, ready-to-use (Roche) and detected with a Chemiluminescence Imaging – Fusion FX SPECTRA instrument (Vilber Lourmat, Collégien, France).

### *In vitro* transcription and confocal laser scanning microscopy

For *in vitro* dsRNA transcription, genomic DNA was isolated from mycelium of *A. fumigatus* CEA17Δ*akuB^KU80^* grown overnight in liquid cultures and used as template for the amplification of 500 base pairs of the *pabA* sequence by PCR. The PCR reaction mix consisted of 200-300 ng template DNA, 10 pmol of each primer and 25 μl of 2x Phusion Flash High-Fidelity PCR Master Mix (Thermo Fisher Scientific). Both, the forward and the reverse primer were designed to contain a T7 promotor at their 5’ end for subsequent dsRNA transcription **(Table S2;** T7_pabA_500_1_for/T7_pab_A500_1_rev). PCR products were loaded onto a 1% agarose gel to check for the correct size and purified with the GeneJET PCR Purification kit (Thermo Fisher Scientific). The products were then used as template for *in vitro* dsRNA transcription using the MEGAscript RNAi Kit (Invitrogen) according to the manufacturer’s protocol with slight modifications. Briefly, the transcription reaction was assembled as specified with the addition of 0.5 μl fluorescein-labelled-UTP Solution (Sigma-Aldrich). The reaction was incubated at 37°C for 4 h and the RNA products were annealed by heating the reaction to 75°C for 5 min and subsequently left to cool at room temperature. Digestion of DNA and single-stranded RNA and the purification of dsRNA was performed as described in the protocol with the exception that the purified dsRNA was eluted in two steps using a total of 100 μL of 95°C ultrapure H_2_O. Quantity and quality of the dsRNA was assessed by fluorometric analysis using a Qubit 4 Fluorometer (Thermo Fisher Scientific). dsRNA was stored in the dark at −80°C until usage.

For fluorescence microscopy, 1×10^4^ freshly collected *A. fumigatus* AfS150 conidia (Lother et al. 2014) were seeded into each well of a 8-well chamber slide (μ-Slide 8 Well, Ibidi) in a total of 100 μl RPMI medium (Sigma-Aldrich), treated with 2 μg of labelled dsRNA, and incubated at 37°C and 5% CO_2_ (v/v) for 16 h. The following day, old media was removed from the samples and fresh media containing calcofluor white was added to the fungus. After 30 min of incubation at room temperature in the dark, hyphae were carefully washed with sterile phosphate-buffered saline (PBS). 100 μl RPMI was added to each chamber and samples were observed by confocal laser scanning microscopy with a Zeiss LSM 780 microscope (Carl Zeiss). Images were processed using the ZEN blue edition software (version 3.4, Carl Zeiss).

### RNA isolation

Fungal mycelium was collected from liquid culture using Miracloth (Millipore) and disrupted in liquid nitrogen using a precooled mortar and pestle. Roughly 0.5 g of homogenized mycelium was transferred into a 2-ml Eppendorf tube. 800 μl TRIzol was added to the ground mycelia and vortexed vigorously. Tubes were frozen briefly for 5 sec in liquid nitrogen and allowed to thaw on ice. 160 μl chloroform was added to the thawed mixture, vortexed, and centrifugated for 5 min at 4°C at full speed. The aqueous upper phase was transferred to a fresh 2-ml tube without disturbing the interphase. RNA extraction from aqueous phase was done with 1 volume of phenol/chloroform/isoamyl alcohol (25:24:1, v/v). Brief vortexing preceded centrifugation for 5 min at 4°C. Extraction was repeated until no more interphase was observable, and this was followed by another extraction with 400 μl chloroform. RNA was precipitated using 400 μl isopropanol for 20 min, followed by pelleting by centrifugation for 20 min at 4°C. The pellet was washed with 700 μl 70% ethanol and air dried at 37°C for 5 min prior to resuspension in RNase free water. The RNA isolation was followed by a DNase treatment using 2 units of TURBO DNase (Thermo Fisher) per 10 μg RNA for 30 min at 37°C in 100 μl total volume. Total RNA was collected using the RNA Clean and Concentrator-25 kit (Zymo Research) according to the manufacturer’s instructions.

### Quantitative RT-PCR (qRT-PCR)

RNA was isolated from cultures of CEA17Δ*akuB^KU80^, pabA*-IRT, and *pksP*-IRT with and without doxycycline induction after 24 h. Complementary DNA (cDNA) was synthesized from RNA according to the iScript cDNA Synthesis Kit protocol (Bio-Rad Laboratories Inc.; Hercules, USA), and the concentration was determined using a Qubit flex fluorometer (Thermo Fisher Scientific). A 20-μl reaction volume per well was prepared in 96-well plate. The PCR reaction mix consisted of: 10 ng cDNA, 4 pmol of each primer pair, 1 μl of 20x Eva Green (Biotium; Hayward, USA) and 10 μl of 2x MyTaq HS Mix (Bioline). Specific primers (qRT_coxV_for / qRT_coxV_rev) for *A. fumigatus* cytochrome C oxidase subunit V (AFUA_5G10560) were used as internal control to normalize the expression of target genes. The cDNAs from four biological samples were used for analysis, and all the reactions were run in triplicate. The reaction was carried out using the QuantStudio 3 Real Time PCR System (Thermo Fisher Scientific) with the following thermal cycling profile: 95°C for 20 s, followed by 40 cycles of amplification (95°C for 5 s, 60°C for 34 s). Relative expression was calculated according to the 2^-ΔΔC^_T_ method (Livak and Schmittgen 2001).

### mRNA Seq and analysis

Total RNA was isolated from three biological replicates of CEA17Δ*akuB^KU80^*, Δ*ppdA/B*, Δ*dclA/B*, and Δ*rrpA/B* knockout strains from conidia, 24-h-, and 48-h-old mycelium. RNA was isolated from mycelia as described above. For *A. fumigatus* conidia, the spores were first homogenized in a bead beater (0.5 mm beads; FastPrep-24, MP Biomedicals) in Trizol before proceeding with the first chloroform extraction step as described in the RNA isolation method above. The RNA isolation was followed by a DNase treatment using 2 units of TURBO DNase (Thermo Fisher) per 10 μg RNA for 30 min at 37°C in 100 μl total volume. Total RNA was then collected using the RNA Clean and Concentrator-25 kit (Zymo Research) according to the manufacturer’s instructions. Total RNA collected were then sent for sequencing. Poly-(A) selection and directional sequencing of mRNA was performed by Novogene using paired-end 150-bp read sequencing on a NovaSeq 6000.

Preprocessing of raw reads including quality control and gene abundance estimation was done with the GEO2RNaseq pipeline (v0.9.12; (Seelbinder et al. 2019)) in R (version 3.5.1). Quality analysis was done with FastQC (v0.11.8) before and after trimming. Read-quality trimming was done with Trimmomatic (v0.36). Reads were rRNA-filtered using SortMeRNA (v2.1) with a single rRNA database combining all rRNA databases shipped with SortMeRNA. Reference annotation was created by extracting and combining exon features from corresponding annotation files. Reads were mapped against the reference genome of *A. fumigatus* (Af293, ASM265v1) using HiSat2 (v2.1.0, paired-end mode). Gene abundance estimation was done with featureCounts (v1.28.0) in paired-end mode with default parameters. MultiQC version 1.7 was finally used to summarize and assess the quality of the output of FastQC, Trimmomatic, HiSat, featureCounts and SAMtools. The count matrix with gene abundance data without and with median-of-ratios normalization (MRN; (Anders and Huber 2010)) were extracted. Raw files are accessible under the Gene Expression Omnibus accession number GSE223618.

Differential gene expression was analyzed using GEO2RNaseq. Pairwise tests were performed using four statistical tools (DESeq v1.30.0, DESeq2 v1.18.1, limma voom v3.34.6 and edgeR v3.20.7) to report *P*-values and multiple testing corrected P-values using the false-discovery rate method q = FDR(p) for each tool. Gene expression differences were considered significant if they were reported significant by all four tools (q ≤ 0.05 and |log_2_ (MRN)| ≤ 1). GO-Slim analysis was performed on genes identified as significantly differentially expressed by all four methods described above using https://fungidb.org (Amos et al. 2022) with a P-Value cutoff of 0.05 and limiting the search to only GO-Slim terms.

### Proteome analysis

Spores were harvest from 5-day old AMM agar plates incubated at 37°C. Spores were pelleted in 2-ml screw cap tubes and a 300 μl equivalent of 0.5 mm beads were added. Bead beating was done in lysis buffer (1% (w/v) SDS, 150 mM NaCl, 100 mM TEAB (triethyl ammonium bicarbonate), with the addition of each 1 tablet cOmplete ULTRA protease inhibitor cocktail and PhosStop (both Roche) per 10 ml lysis buffer), 3 times for 20 secs with 2-minute rest on ice in between. Harvested mycelium from 48-h cultures was disrupted by mortar and pestle with liquid nitrogen. Cell debris was then homogenized in lysis buffer. All samples were then incubated at 37°C in a water bath sonicator for 30 min with 0.5 μl Benzonase nuclease (250 U/μl). Proteins were separated from insolubilized debris by centrifugation (15 min, 18000 × g). Each 100 μg of total protein per sample was diluted with 100 mM TEAB to gain a final volume of 100 μl. Subsequently, cysteine thiols were reduced and carbamidomethylated for 30 min at 70°C by addition of each 2 μl of 500 mM TCEP (tris(2-carboxyethyl)phosphine) and 625 mM 2-chloroacetamide (CAA). The samples were further cleaned up by methanol-chloroform-water precipitation using the protocol of Wessel and Flügge (Wessel and Flugge 1984). Protein precipitates were resolubilized in 5% trifluoroethanol of aqueous 100 mM TEAB and digested for 18 h with Trypsin+LysC (Promega) at a ratio of 25:1 protein:protease. Samples were evaporated in a vacuum concentrator (Eppendorf). Tryptic peptides were resolubilized in 30 μl of 0.05% TFA and 2% ACN in H_2_O filtered through 10 kDa MWCO PES membrane spin filters (VWR) at 14,000×*g* for 15 min and transferred to HPLC vials.

### LC-MS/MS analysis and protein database search

An improvement in technology was implemented between the two LC-MS/MS proteomics experiments described in this manuscript. Each method is described in detail below for clarity. Method 1 (M1) is related to 48-h mycelial proteomics (3 biological replicates) and method 2 (M2) to conidial proteomics (4 biological replicates). Each biological replicate was measured in triplicate (3 analytical replicates). LC-MS/MS analysis was performed on an Ultimate 3000 nano RSLC system connected to either a QExactive Plus (M1) or an Orbitrap Exploris 480 mass spectrometer (both Thermo Fisher Scientific, Waltham, MA, USA) equipped with a FAIMS ion mobility interface (M2). Peptide trapping for 5 min on an Acclaim Pep Map 100 column (2 cm x 75 μm, 3 μm) at 5 μL/min was followed by separation on an analytical Acclaim Pep Map RSLC nano column (50 cm x 75 μm, 2μm). Mobile phase gradient elution of eluent A (0.1% (v/v) formic acid in water) mixed with eluent B (0.1% (v/v) formic acid in 90/10 acetonitrile/water) was performed using the following gradients. M1: 0–5 min at 4% B, 30 min at 7% B, 60 min at 10% B, 100 min at 15% B, 140 min at 25% B, 180 min at 45% B, 200 min at 65%, 210–215 min at 96% B, 215.1–240 min at 4% B. M2: 0 min at 4% B, 20 min at 6% B, 45 min at 10% B, 75 min at 16% B, 105 min at 25% B, 135 min at 45% B, 150 min at 65% B, 160-165 min at 96% B, 165.1-180 min at 4% B. Positively charged ions were generated at spray voltage of 2.2 kV using a stainless steel emitter attached to the Nanospray Flex Ion Source (Thermo Fisher Scientific). The quadrupole/orbitrap instrument was operated in Full MS / data-dependent MS2 mode to trigger fragmentation of the Top 15 precursor ions (M1). For M2 the number of data-dependent scans was related to the cycle time of 1.5 s (M2). Precursor ions were monitored at *m/z* 300-1500 (M1) or *m/z* 300-1200 (M2) at a resolution of 140,000 (M1) or 120,000 (M2) FWHM (full width at half maximum) using a maximum injection time (ITmax) of 120 ms (M1) or 50 ms (M2) and an AGC (automatic gain control) target of 3×10^6^ (M1) and a normalized AGC target of 300% (M2), respectively. Precursor ions with a charge state of z=2-5 were filtered at an isolation width of *m/z* 1.6 (M1) or *m/z* 4.0 (M2) for further fragmentation at 30% (M1) or 28% (M2) HCD collision energy. MS2 ions were scanned at 17,500 FWHM (ITmax=120 ms, AGC= 2×10^5^) for M1 and 15,000 FWHM (ITmax=40 ms, AGC= 200%) for M2. 3 different compensation voltages were applied (−48V, −63V, −78V) for the FAIMS interface (M2).

Tandem mass spectra were searched against the FungiDB database (https://fungidb.org/common/downloads/Current_Release/AfumigatusAf293/fasta/data/FungiDB-57_AfumigatusAf293_AnnotatedProteins.fasta) of *Aspergillus fumigatus* Af293 (2021/07/09; YYYY/MM/DD) for M1 and against the UniProt pan proteome databases (2022/12/21 (YYYY/MM/DD) of *Neosartorya fumigata / Aspergillus fumigatus* (https://ftp.uniprot.org/pub/databases/uniprot/current_release/knowledgebase/pan_proteomes/UP000002530.fasta.gz) for M2 using Thermo Proteome Discoverer (PD) 2.4 (M1) or 3.0 (M2). The following database search engines and respective scoring thresholds (in parenthesis) have been applied for the different methods. M1: Mascot 2.4.1 (>30), Sequest HT (>4), MS Amanda 2.0 (>300), and MS Fragger 3.2 (>8). M2: Mascot 2.8 (>30), Comet (>3), MS Amanda 2.0 (>300), Sequest HT with and without INFERYS Rescoring (>3), and CHIMERYS (>2). Two missed cleavages were allowed for the tryptic digestion. The precursor mass tolerance was set to 10 ppm and the fragment mass tolerance was set to 0.02 Da. Modifications were defined as dynamic Met oxidation, phosphorylation of Ser, Thr, and Tyr (M2), protein N-term acetylation with and without Met-loss as well as static Cys carbamidomethylation. A strict false discovery rate (FDR) < 1% (peptide and protein level) and aforementioned search engine score threshold values were required for positive protein hits. The Percolator node and a reverse decoy database was used for q-value validation of spectral matches. Only rank 1 proteins and peptides of the top scored proteins were counted. Label-free protein quantification was based on the Minora algorithm using the precursor abundance based on intensity and a signal-to-noise ratio >5. Normalization was performed by using the total peptide amount method. Imputation of missing quan values was applied by using abundance values of 75% of the lowest abundance identified. Differential protein abundance was defined as a fold change of > 2, *P*-value/ABS(log_4_ratio) ≤ 0.05 and at least identified in 2 of 3 (M1) or 3 of 4 replicates (M2) of the sample group with the highest abundance **(Dataset S4-5).** Heatmaps were created in R v. 4.2.2 using pheatmap v. 1.0.12 by first Log_2_-transforming normalized abundance data and then calculating Z scores for proteins that exhibited at least a log_2_(FC)=2 and a P-value ≤ 0.05 in one of the conditions.

### Library preparation for small RNA-sequencing and bioinformatic analysis

Total RNA was isolated from three separate biological experiments using *A. fumigatus* CEA10 conidia (5 days on AMM agar plates at 37°C) and mycelium after 24 and 48 h of growth in AMM liquid culture at 37°C using the Universal RNA Kit (Roboklon) with slight modifications. Namely, conidia and mycelia at different time points were harvested and resuspended in RL buffer followed by bead beating 3x for 30 s each using 0.5 mm glass beads (Carl Roth). The total RNA was then isolated according to the yeast protocol provided by the manufacturer (Roboklon) using an on-column DNase treatment (Lucigen Baseline-ZERO DNase). The RNA isolation was followed by an additional DNase treatment using 4 units of TURBO DNase (Thermo Fisher) for 30 min at 37°C in 100 μl total volume. The small RNAs were then collected using the RNA Clean and Concentrator-25 kit (Zymo Research) according to the manufacturer’s instructions for small RNA enrichment.

Since the 5’ end of *A. fumigatus* small RNAs are not well-defined, the isolated RNA was treated with 1 μl (5 units) RppH enzyme (New England Biolabs) in NEB Buffer 2 (50 mM NaCl, 10 mM Tris-HCl, 10 mM MgCl_2_, and 1 mM DTT at pH 7.9) for 1 h at 37°C to facilitate removal of a pyrophosphate from the 5’ end of triphosphorylated RNA. The reaction was ended with EDTA for 5 min at 65°C followed by cleanup using the RNA Clean and Concentrator-25 kit (Zymo Research) as described above. RNA was quantified using the Qubit 4 Fluorometer (Thermo Fisher). Small RNA libraries were then prepared from 10 ng of small RNA using the NEBNext Multiplex Small RNA Library Prep Set for Illumina (New England Biolabs #E7300S) according to the manufacturer’s recommendation using NEBNext Multiplex Oligos for Illumina (Index Primer Set 1, #E7335S; **Table S1).** Size selection was performed according to the manufacturer using AMPure XP beads (Beckman Coulter). Libraries were validated for the correct size by polyacrylamide gel electrophoresis. The barcoded libraries were pooled without further adjustment, followed by 50-cycle-single-end sequencing using the Illumina HiSeq 4000 platform (Eurofins Genomics). In this case, the sRNA data produced are not strictly suitable for comparison of read abundance between time points but were deemed appropriate for sRNA discovery.

Adaptors were trimmed from sRNA sequencing data using BBDuk from BBTools v38.22 (https://sourceforge.net/projects/bbmap/) and quality profiles assessed using FastQC. Candidate sRNAs were identified from merged replicate data using Manatee v1.2 (Handzlik et al. 2020) along with the Ensembl *A. fumigatus* A1163 genome (ASM15014v1) sequence and annotated transfer RNAs as a reference. Sequence hits between 19 and 25 nucleotides in length and with more than 200 counts were retained for further analysis. Hits were visually inspected using sRNA- and mRNA-seq data and the complete *A. fumigatus* gene annotation viewed in Integrative Genomics Viewer (IGV) to validate their existence and identify antisense overlap with gene transcription (Thorvaldsdottir et al. 2013). FASTA sequences for the final candidate sRNAs were extracted using gffread from GFF utilities (Pertea and Pertea 2020).

## Data availability

The mass spectrometry proteomics data were deposited in the ProteomeXchange Consortium via the PRIDE partner repository (Perez-Riverol et al. 2022) with the dataset identifiers PXD033984 (48-h mycelium) and PXD039739 (conidia). Small RNA- and mRNA-sequencing datasets can be found on GEO under identifiers GSE223619 and GSE223618, respectively.

## Statistical Analysis

The proteomics data were analyzed as described. For the remaining analyses, *P*-values were determined by ANOVA or two-tailed Student’s t-tests where appropriate using Prism 5.0.0 software (GraphPad Software). Differences between the groups were considered significant at a *P*-value of <0.05. Throughout the article, significance is denoted as follows: *, *P* ≤ 0.05; **, *P* ≤ 0.01, ***, *P* ≤ 0.001; ns, nonsignificant.

## ACKNOWLEDGEMENTS

We would like to thank Pamela Lehenberger, Bhawana Israni, Carmen Schult, and Laura Broschat for excellent technical assistance. The work presented here was generously supported by the Federal Ministry for Education and Research (BMBF: https://www.bmbf.de/), Germany, Project FKZ 01K12012 “RFIN – RNA-Biologie von Pilzinfektionen”. Additional funding support came from the Deutsche Forschungsgemeinschaft (DFG; German Research Foundation) under Germany’s Excellence Strategy – EXC 2051 – Project-ID 390713860 and the DFG-funded Collaborative Research Center/Transregio FungiNet 124 ‘Pathogenic fungi and their human host: Networks of Interaction’ (210879364, project A1, INF, and Z2). The authors declare no conflicts of interest.

## SUPPLEMENTARY FIGURES

**FIG S1. The RNAi machinery incurs only minor-to-moderate genetic variation.**

Histogram depicting the number of variants and their functional impact in RNAi genes from 300 of *A. fumigatus* genomes. Genetic changes are reported relative to the Af293 reference genome. DNA base changes resulting in synonymous substitutions or those in intergenic regions are not shown. Full summary in **Dataset S1.**

**FIG S2. Southern blot validation of knockout strains.**

Southern blot images showing confirmation of gene deletions in knockout strains. Numbers indicate candidate knockout strains collected from transformation plates. Candidates were digested with appropriate restriction enzymes to produce fragments of differing sizes in the wild type (WT) and knockout for each gene. One of the flanking regions was used as the probe by using primers from the creation of the knockout as well as the addition of DIG-UTP to the PCR reaction. Stars indicate strains with construct of anticipated size.

**FIG S3. RNAi does not contribute to stress response or drug resistance.**

**(A)** Droplet stress assays for wild-type (WT), Δ*dclA/B*, Δ*ppdA/B*, and Δ*rrpA/B* strains grown on AMM agar plates supplemented with indicated stress inducers for 48 h. Wild type and RNAi double knockouts were grown on increasing concentrations of (B) voriconazole and (C) tebuconazole in AMM media. The minimum inhibitory concentration (MIC) was determined after 48 h. Images shown for each stress condition in A-C) are representative of three independent experiments.

**FIG S4. The *A. fumigatus* cell wall limits uptake of long double-stranded RNA.**

**(A)** The *A. fumigatus* cell wall limits the delivery of extracellular RNA as shown by confocal laser scanning microscopy of calcofluor white-stained (blue) *A. fumigatus* AfS150 hyphae expressing a cytoplasmic tdTomato (red), which were incubated with *in vitro*-transcribed fluorescein-UTP labelled dsRNA (green). (B) Schematic diagram of the constructs (pSilent-*pksP1*, pSilent-*pksP2*, and pSilent-*pabA*) used to induce RNAi. Sense and anti-sense *pksP* PCR product **(green)** were placed downstream of an inducible Tet-On promoter **(blue).** The inverted-repeat region and antifungal selection genes **(*hph*** and ***ble*)** for each plasmid were collectively flanked by 5’ and 3’ regions **(black)** amplified from template of a non-functional *pyrG* locus for homologous recombination.

**FIG S5. Principle component analysis and hierarchical clustering of mRNA-seq analysis.**

(A) Principle component analysis performed on MRN-normalized read counts showing clear separation of samples based on timepoint. (B) Hierarchical clustering of mRNA-seq samples, revealing close grouping of the samples based on time point.

**FIG S6. GO-slim analysis shows links to RNA binding and the nucleolus for RNAi components.**

GO-Slim analysis (using FungiDB.org) for Cellular Component and Molecular Function generated using differentially expressed genes from conidia **(Fig. 3A)** as input gene sets. Left panels show categories for up- or down-regulated genes. Right panels show total number of genes that were found in the particular category.

**FIG S7. Proteomics reveals altered protein abundance in double knockouts in 48-h mycelium.**

(A) Clustered heatmap depicting Z-scored normalized average abundance data of 48-h mycelium proteomics samples for double knockouts and wild type. Included are cases where proteins exhibited at least a log_2_(FC)=2 and a P-value ≤ 0.05 in one of the conditions. Normalized abundances were determined using area under the curve from label-free quantification. (B-D) Volcano plots showing expression ratio for double knockout compared to wild type for (B) Δ*dclA/B*, **(C)** Δ*ppdA/B*, and (D) Δ*rrpA/B*. The y-axis shows the −log_10_ transformed P-value, where higher values are more significant. Orange circles show proteins that are at least 2-fold changed and with a P-value ≤ 0.05.

**FIG S8. Images of fungal strains evolved over six generations without selective pressure.**

Double knockout strains were grown as 3 biological lineages and passaged for 6 generations on AMM agar plates. Each generation was allowed to grow at 37°C for 7 days before the next passage. Each plate for each lineage was photographed before harvesting spores. The slight color changes are due to changes in ambient light during image acquisition. The * indicates a second evolution experiment performed at a later date to minimize cross-contamination events.

## SUPPLEMENTARY TABLES

**Table S1.** Statistics for sRNA-seq analysis.

| Condition | Replicate | Barcode | Raw Reads |
| --- | --- | --- | --- |
| Conidia | 1 | ATCACG | 8,846,270 |
| Conidia | 2 | CGATGT | 3,733,570 |
| Conidia | 3 | TGACCA | 17,100,200 |
| 24 h Mycelium | 1 | ACAGTG | 15,884,500 |
| 24 h Mycelium | 2 | GCCAAT | 12,093,800 |
| 24 h Mycelium | 3 | ACTTGA | 1,282,660 |
| 48 h Mycelium | 1 | GATCAG | 37,359,400 |
| 48 h Mycelium | 2 | GGCTAC | 96,662,100 |
| 48 h Mycelium | 3 | CTTGTA | 85,428,800 |

**Table S2.**
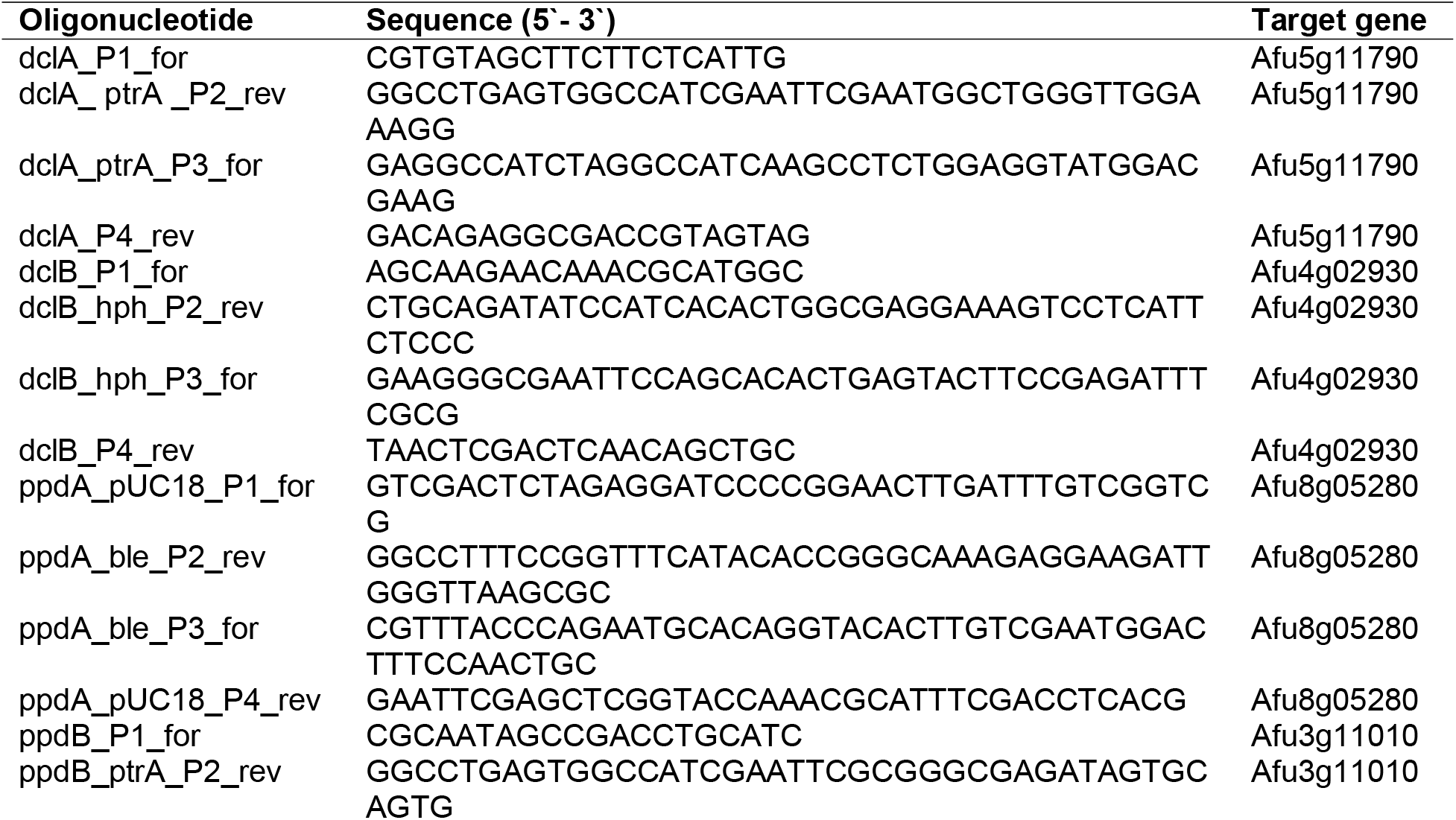
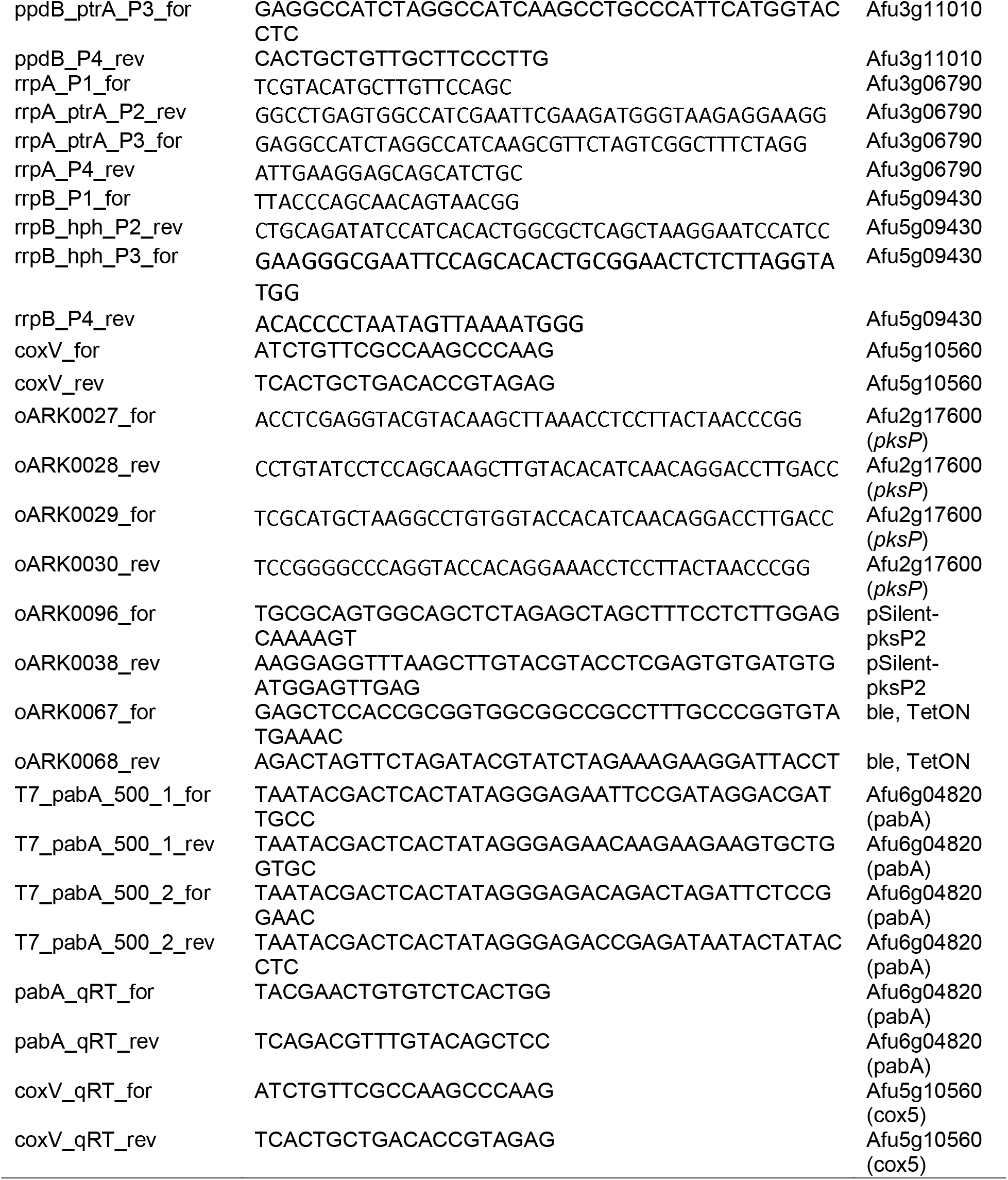
Oligonucleotides used in this study.

**Table S3.**
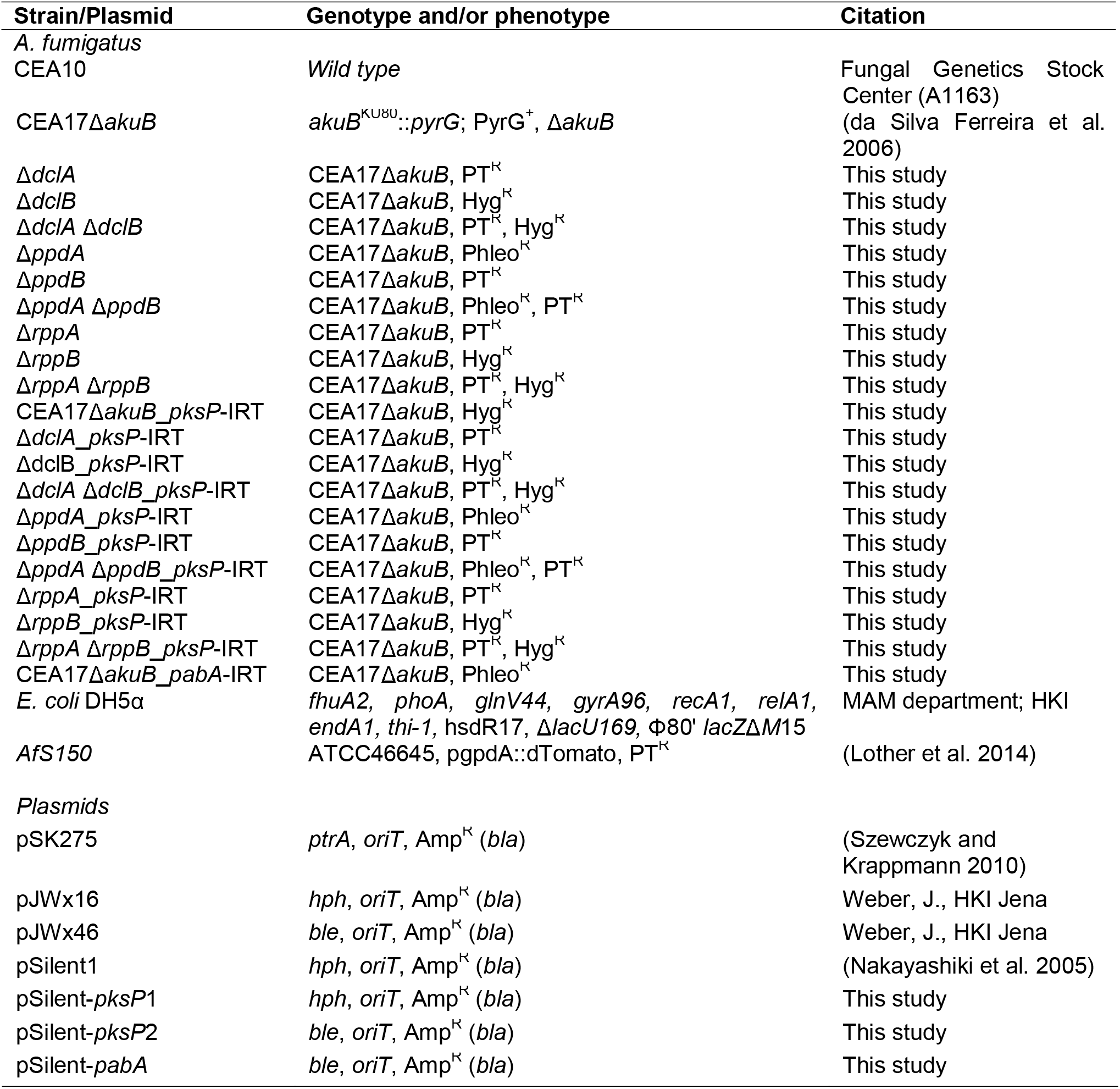
Strains and plasmids used in this study.

## SUPPLEMENTARY DATASETS

**Dataset S1.** List of the genetic variants identified in RNAi machinery of 300 sequenced *A. fumigatus* isolates first described in (Barber et al. 2021).

**Dataset S2.** Normalized read counts generated by mRNA-seq of RNAi double knockouts.

**Dataset S3.** GO-slim analysis of genes differentially expressed in conidia of RNAi double knockouts.

**Dataset S4.** LC-MS/MS proteomics results performed on conidia of wild-type and RNAi double knockout strains.

**Dataset S5.** LC-MS/MS proteomics results performed on 48-hour-old mycelia of wild-type and RNAi double knockout strains.

**Dataset S6.** Small RNAs identified in this study.

## REFERENCES

Abdel-Hadi AM, Caley DP, Carter DR, Magan N. 2011. Control of aflatoxin production of *Aspergillus flavus* and *Aspergillus parasiticus* using RNA silencing technology by targeting *aflD* (*nor-1*) gene. Toxins (Basel) 3: 647–659.

Agrawal N, Dasaradhi PV, Mohmmed A, Malhotra P, Bhatnagar RK, Mukherjee SK. 2003. RNA interference: biology, mechanism, and applications. Microbiol Mol Biol Rev 67: 657–685.

Alexander WG, Raju NB, Xiao H, Hammond TM, Perdue TD, Metzenberg RL, Pukkila PJ, Shiu PK. 2008. DCL-1 colocalizes with other components of the MSUD machinery and is required for silencing. Fungal Genet Biol 45: 719–727.

Amos B, Aurrecoechea C, Barba M, Barreto A, Basenko EY, Bazant W, Belnap R, Blevins AS, Bohme U, Brestelli J et al. 2022. VEuPathDB: the eukaryotic pathogen, vector and host bioinformatics resource center. Nucleic Acids Res 50: D898–D911.

Anders S, Huber W. 2010. Differential expression analysis for sequence count data. Genome Biol 11: R106.

Barber AE, Riedel J, Sae-Ong T, Kang K, Brabetz W, Panagiotou G, Deising HB, Kurzai O. 2020. Effects of Agricultural Fungicide Use on *Aspergillus fumigatus* Abundance, Antifungal Susceptibility, and Population Structure. mBio 11.

Barber AE, Sae-Ong T, Kang K, Seelbinder B, Li J, Walther G, Panagiotou G, Kurzai O. 2021. *Aspergillus fumigatus* pan-genome analysis identifies genetic variants associated with human infection. Nat Microbiol 6: 1526–1536.

Beauvais A, Bozza S, Kniemeyer O, Formosa C, Balloy V, Henry C, Roberson RW, Dague E, Chignard M, Brakhage AA et al. 2013. Deletion of the alpha-(1,3)-glucan synthase genes induces a restructuring of the conidial cell wall responsible for the avirulence *of Aspergillus fumigatus*. PLoS Pathog 9: e1003716.

Bernstein DA, Vyas VK, Weinberg DE, Drinnenberg IA, Bartel DP, Fink GR. 2012. *Candida albicans* Dicer (CaDcr1) is required for efficient ribosomal and spliceosomal RNA maturation. Proc Natl Acad Sci U S A 109: 523–528.

Billmyre RB, Calo S, Feretzaki M, Wang X, Heitman J. 2013. RNAi function, diversity, and loss in the fungal kingdom. Chromosome Res 21: 561–572.

Bostian KA, Hopper JE, Rogers DT, Tipper DJ. 1980. Translational analysis of the killer-associated virus-like particle dsRNA genome of S. *cerevisiae:* M dsRNA encodes toxin. Cell 19: 403–414.

Brennan P. 2018. drawProteins: a Bioconductor/R package for reproducible and programmatic generation of protein schematics. F1000Res 7: 1105.

Brown JS, Aufauvre-Brown A, Brown J, Jennings JM, Arst H, Jr., Holden DW. 2000. Signature-tagged and directed mutagenesis identify PABA synthetase as essential for *Aspergillus fumigatus* pathogenicity. Mol Microbiol 36: 1371–1380.

Bruch A, Kelani AA, Blango MG. 2021. RNA-based therapeutics to treat human fungal infections. Trends Microbiol.

Cai Q, He B, Weiberg A, Buck AH, Jin H. 2019. Small RNAs and extracellular vesicles: New mechanisms of cross-species communication and innovative tools for disease control. PLoS Pathog 15: e1008090.

Calo S, Nicolas FE, Lee SC, Vila A, Cervantes M, Torres-Martinez S, Ruiz-Vazquez RM, Cardenas ME, Heitman J. 2017. A non-canonical RNA degradation pathway suppresses RNAi-dependent epimutations in the human fungal pathogen *Mucor circinelloides*. PLoS Genet 13: e1006686.

Carreras-Villasenor N, Esquivel-Naranjo EU, Villalobos-Escobedo JM, Abreu-Goodger C, Herrera-Estrella A. 2013. The RNAi machinery regulates growth and development in the filamentous fungus *Trichoderma atroviride*. Mol Microbiol 89: 96–112.

Chen Y, Gao Q, Huang M, Liu Y, Liu Z, Liu X, Ma Z. 2015. Characterization of RNA silencing components in the plant pathogenic fungus *Fusarium graminearum*. Sci Rep 5: 12500.

D’Souza CA, Kronstad JW, Taylor G, Warren R, Yuen M, Hu G, Jung WH, Sham A, Kidd SE, Tangen K et al. 2011. *Genome variation in Cryptococcus gattii*, an emerging pathogen of immunocompetent hosts. mBio 2: e00342-00310.

da Silva Ferreira ME, Kress MR, Savoldi M, Goldman MH, Hartl A, Heinekamp T, Brakhage AA, Goldman GH. 2006. The akuB(KU80) mutant deficient for nonhomologous end joining is a powerful tool for analyzing pathogenicity in *Aspergillus fumigatus*. Eukaryot Cell 5: 207–211.

Dang Y, Yang Q, Xue Z, Liu Y. 2011. RNA interference in fungi: pathways, functions, and applications. Eukaryot Cell 10: 1148–1155.

Ender C, Meister G. 2010. Argonaute proteins at a glance. J Cell Sci 123: 1819–1823.

Gaffar FY, Imani J, Karlovsky P, Koch A, Kogel KH. 2019. Different Components of the RNA Interference Machinery Are Required for Conidiation, Ascosporogenesis, Virulence, Deoxynivalenol Production, and Fungal Inhibition by Exogenous Double-Stranded RNA in the Head Blight Pathogen *Fusarium graminearum*. Front Microbiol 10: 1662.

Gibbons JG, D’Avino P, Zhao S, Cox GW, Rinker DC, Fortwendel JR, Latge J-P. 2022. Comparative Genomics Reveals a Single Nucleotide Deletion in *pksP* That Results in White-Spore Phenotype in Natural Variants of *Aspergillus fumigatus*. Frontiers in Fungal Biology 3.

Gusa A, Williams JD, Cho JE, Averette AF, Sun S, Shouse EM, Heitman J, Alspaugh JA, Jinks-Robertson S. 2020. Transposon mobilization in the human fungal pathogen *Cryptococcus* is mutagenic during infection and promotes drug resistance in vitro. Proc Natl Acad Sci U S A 117: 9973–9980.

Hall IM, Noma K, Grewal SI. 2003. RNA interference machinery regulates chromosome dynamics during mitosis and meiosis in fission yeast. Proc Natl Acad Sci U S A 100: 193–198.

Hammond TM, Andrewski MD, Roossinck MJ, Keller NP. 2008a. *Aspergillus* mycoviruses are targets and suppressors of RNA silencing. Eukaryot Cell 7: 350–357.

Hammond TM, Bok JW, Andrewski MD, Reyes-Dominguez Y, Scazzocchio C, Keller NP. 2008b. RNA silencing gene truncation in the filamentous fungus *Aspergillus nidulans*. Eukaryot Cell 7: 339–349.

Hammond TM, Keller NP. 2005. RNA silencing in *Aspergillus nidulans* is independent of RNA-dependent RNA polymerases. Genetics 169: 607–617.

Handzlik JE, Tastsoglou S, Vlachos IS, Hatzigeorgiou AG. 2020. Manatee: detection and quantification of small non-coding RNAs from next-generation sequencing data. Sci Rep 10: 705.

Hutvagner G, Simard MJ. 2008. Argonaute proteins: key players in RNA silencing. Nat Rev Mol Cell Biol 9: 22–32.

Janbon G, Maeng S, Yang DH, Ko YJ, Jung KW, Moyrand F, Floyd A, Heitman J, Bahn YS. 2010. Characterizing the role of RNA silencing components in *Cryptococcus neoformans*. Fungal Genet Biol 47: 1070–1080.

Jinek M, Doudna JA. 2009. A three-dimensional view of the molecular machinery of RNA interference. Nature 457: 405–412.

Jochl C, Loh E, Ploner A, Haas H, Huttenhofer A. 2009. Development-dependent scavenging of nucleic acids in the filamentous fungus *Aspergillus fumigatus*. RNA Biol 6: 179–186.

Kadotani N, Nakayashiki H, Tosa Y, Mayama S. 2003. RNA silencing in the phytopathogenic fungus *Magnaporthe oryzae*. Mol Plant Microbe Interact 16: 769–776.

Kalleda N, Naorem A, Manchikatla RV. 2013. Targeting fungal genes by diced siRNAs: a rapid tool to decipher gene function in *Aspergillus nidulans*. PLoS One 8: e75443.

Ketting RF. 2011. The many faces of RNAi. Dev Cell 20: 148–161.

Khatri M, Rajam MV. 2007. Targeting polyamines of *Aspergillus nidulans* by siRNA specific to fungal ornithine decarboxylase gene. Med Mycol 45: 211–220.

Langfelder K, Jahn B, Gehringer H, Schmidt A, Wanner G, Brakhage AA. 1998. Identification of a polyketide synthase gene (*pksP) of Aspergillus fumigatus* involved in conidial pigment biosynthesis and virulence. Med Microbiol Immunol 187: 79–89.

Latge JP. 1999. *Aspergillus fumigatus* and aspergillosis. Clin Microbiol Rev 12: 310–350.

Livak KJ, Schmittgen TD. 2001. Analysis of relative gene expression data using real-time quantitative PCR and the 2(-Delta Delta C(T)) Method. Methods 25: 402–408.

Lother J, Breitschopf T, Krappmann S, Morton CO, Bouzani M, Kurzai O, Gunzer M, Hasenberg M, Einsele H, Loeffler J. 2014. Human dendritic cell subsets display distinct interactions with the pathogenic mould *Aspergillus fumigatus*. Int J Med Microbiol 304: 1160–1168.

Martienssen R, Moazed D. 2015. RNAi and heterochromatin assembly. Cold Spring Harb Perspect Biol 7: a019323.

Moazed D. 2009. Small RNAs in transcriptional gene silencing and genome defence. Nature 457: 413–420.

Mousavi B, Hedayati MT, Teimoori-Toolabi L, Guillot J, Alizadeh A, Badali H. 2015. cyp51A gene silencing using RNA interference in azole-resistant *Aspergillus fumigatus*. Mycoses 58: 699–706.

Mouyna I, Henry C, Doering TL, Latge JP. 2004. Gene silencing with RNA interference in the human pathogenic fungus *Aspergillus fumigatus*. FEMS Microbiol Lett 237: 317–324.

Mukherjee K, Campos H, Kolaczkowski B. 2013. Evolution of animal and plant dicers: early parallel duplications and recurrent adaptation of antiviral RNA binding in plants. Mol Biol Evol 30: 627–641.

Murata T, Kadotani N, Yamaguchi M, Tosa Y, Mayama S, Nakayashiki H. 2007. siRNA-dependent and - independent post-transcriptional cosuppression of the LTR-retrotransposon MAGGY in the phytopathogenic fungus *Magnaporthe oryzae*. Nucleic Acids Res 35: 5987–5994.

Nakayashiki H, Hanada S, Nguyen BQ, Kadotani N, Tosa Y, Mayama S. 2005. RNA silencing as a tool for exploring gene function in ascomycete fungi. Fungal Genet Biol 42: 275–283.

Nami S, Baradaran B, Mansoori B, Kordbacheh P, Rezaie S, Falahati M, Mohamed Khosroshahi L, Safara M, Zaini F. 2017. The Utilization of RNA Silencing Technology to Mitigate the Voriconazole Resistance of *Aspergillus Flavus;* Lipofectamine-Based Delivery. Adv Pharm Bull 7: 53–59.

Nicolas FE, Garre V. 2016. RNA Interference in Fungi: Retention and Loss. Microbiol Spectr 4.

Nicolas FE, Moxon S, de Haro JP, Calo S, Grigoriev IV, Torres-Martinez S, Moulton V, Ruiz-Vazquez RM, Dalmay T. 2010. Endogenous short RNAs generated by Dicer 2 and RNA-dependent RNA polymerase 1 regulate mRNAs in the basal fungus *Mucor circinelloides*. Nucleic Acids Res 38: 5535–5541.

Nicolas FE, Vila A, Moxon S, Cascales MD, Torres-Martinez S, Ruiz-Vazquez RM, Garre V. 2015. The RNAi machinery controls distinct responses to environmental signals in the basal fungus *Mucor circinelloides*. BMC Genomics 16: 237.

Nuss DL. 2011. Mycoviruses, RNA silencing, and viral RNA recombination. Adv Virus Res 80: 25–48.

Oberegger H, Zadra I, Schoeser M, Haas H. 2000. Iron starvation leads to increased expression of Cu/Zn-superoxide dismutase in *Aspergillus*. FEBS Lett 485: 113–116.

Ozkan S, Mohorianu I, Xu P, Dalmay T, Coutts RHA. 2017. Profile and functional analysis of small RNAs derived from *Aspergillus fumigatus* infected with double-stranded RNA mycoviruses. BMC Genomics 18: 416.

Pascovici D, Handler DC, Wu JX, Haynes PA. 2016. Multiple testing corrections in quantitative proteomics: A useful but blunt tool. Proteomics 16: 2448–2453.

Perez-Riverol Y, Bai J, Bandla C, Garcia-Seisdedos D, Hewapathirana S, Kamatchinathan S, Kundu DJ, Prakash A, Frericks-Zipper A, Eisenacher M et al. 2022. The PRIDE database resources in 2022: a hub for mass spectrometry-based proteomics evidences. Nucleic Acids Res 50: D543–D552.

Pertea G, Pertea M. 2020. GFF Utilities: GffRead and GffCompare. F1000Res 9.

Qin H, Chen F, Huan X, Machida S, Song J, Yuan YA. 2010. Structure of the *Arabidopsis thaliana* DCL4 DUF283 domain reveals a noncanonical double-stranded RNA-binding fold for protein-protein interaction. RNA 16: 474–481.

Roche B, Arcangioli B, Martienssen RA. 2016. RNA interference is essential for cellular quiescence. Science 354.

Romano N, Macino G. 1992. Quelling: transient inactivation of gene expression in *Neurospora crassa* by transformation with homologous sequences. Mol Microbiol 6: 3343–3353.

Ruiz-Vazquez RM, Nicolas FE, Torres-Martinez S, Garre V. 2015. Distinct RNAi Pathways in the Regulation of Physiology and Development in the Fungus *Mucor circinelloides*. Adv Genet 91: 55–102.

Schauer SE, Jacobsen SE, Meinke DW, Ray A. 2002. DICER-LIKE1: blind men and elephants in *Arabidopsis* development. Trends Plant Sci 7: 487–491.

Seelbinder B, Wolf T, Priebe S, McNamara S, Gerber S, Guthke R, Linde J. 2019. GEO2RNAseq: An easy-to-use R pipeline for complete preprocessing of RNA-seq data. bioRxiv.

Shabalina SA, Koonin EV. 2008. Origins and evolution of eukaryotic RNA interference. Trends Ecol Evol 23: 578–587.

Shopova IA, Belyaev I, Dasari P, Jahreis S, Stroe MC, Cseresnyes Z, Zimmermann AK, Medyukhina A, Svensson CM, Kruger T et al. 2020. Human Neutrophils Produce Antifungal Extracellular Vesicles against *Aspergillus fumigatus*. mBio 11.

Song MS, Rossi JJ. 2017. Molecular mechanisms of Dicer: endonuclease and enzymatic activity. Biochem J 474: 1603–1618.

Southern E. 2006. Southern blotting. Nat Protoc 1: 518–525.

Szewczyk E, Krappmann S. 2010. Conserved regulators of mating are essential for *Aspergillus fumigatus* cleistothecium formation. Eukaryot Cell 9: 774–783.

Thorvaldsdottir H, Robinson JT, Mesirov JP. 2013. Integrative Genomics Viewer (IGV): high-performance genomics data visualization and exploration. Brief Bioinform 14: 178–192.

Torres-Martinez S, Ruiz-Vazquez RM. 2017. The RNAi Universe in Fungi: A Varied Landscape of Small RNAs and Biological Functions. Annu Rev Microbiol 71: 371–391.

Torri A, Jaeger J, Pradeu T, Saleh MC. 2022. The origin of RNA interference: Adaptive or neutral evolution? PLoS Biol 20: e3001715.

Wang X, Wang P, Sun S, Darwiche S, Idnurm A, Heitman J. 2012. Transgene induced co-suppression during vegetative growth in *Cryptococcus neoformans*. PLoS Genet 8: e1002885.

Weidner G, d’Enfert C, Koch A, Mol PC, Brakhage AA. 1998. Development of a homologous transformation system for the human pathogenic fungus *Aspergillus fumigatus* based on the pyrG gene encoding orotidine 5’-monophosphate decarboxylase. Curr Genet 33: 378–385.

Wessel D, Flugge UI. 1984. A method for the quantitative recovery of protein in dilute solution in the presence of detergents and lipids. Anal Biochem 138: 141–143.

Xiao H, Alexander WG, Hammond TM, Boone EC, Perdue TD, Pukkila PJ, Shiu PK. 2010. QIP, a protein that converts duplex siRNA into single strands, is required for meiotic silencing by unpaired DNA. Genetics 186: 119–126.

