## Supplementary figures and images for "Disruption of the *Aspergillus fumigatus* RNA interference machinery alters the conidial transcriptome"

### Supplemental Figures

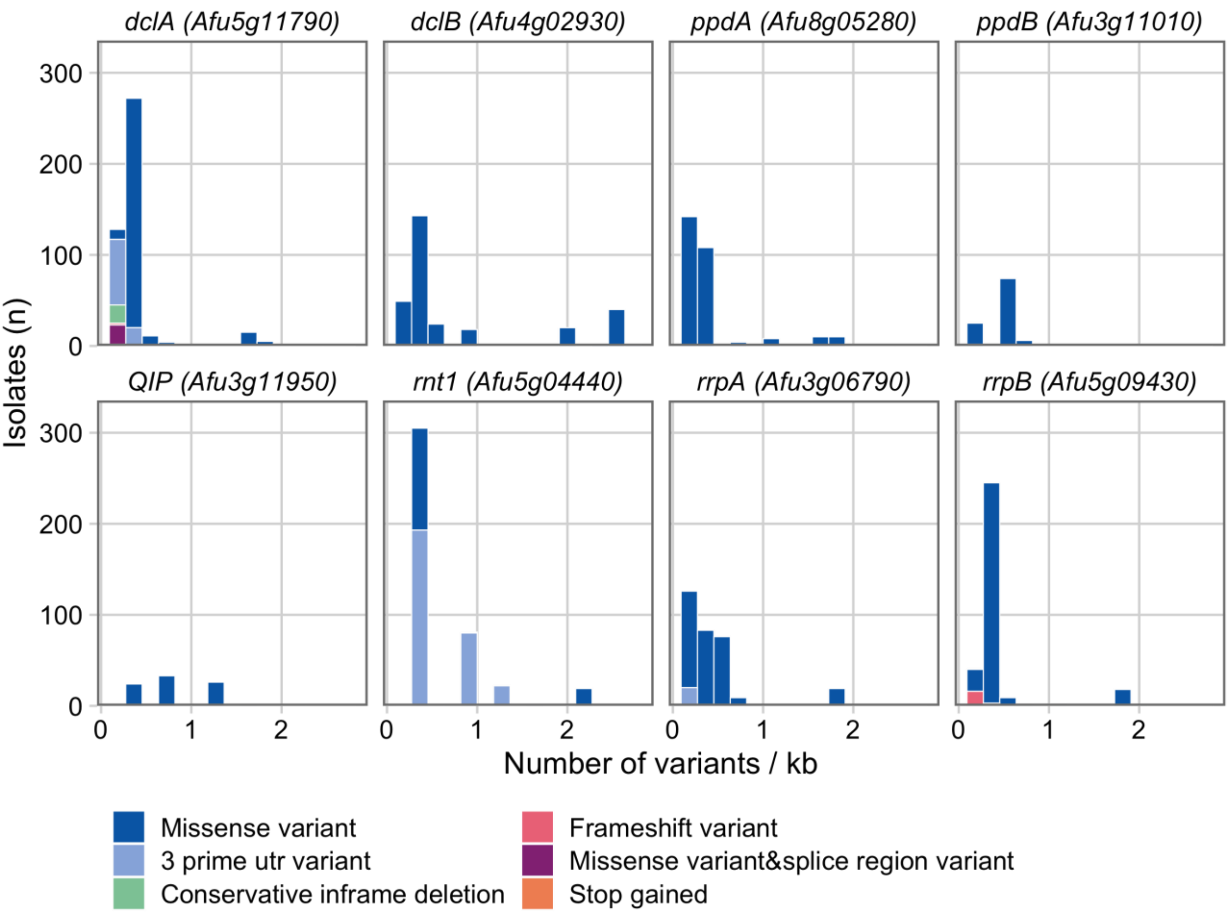

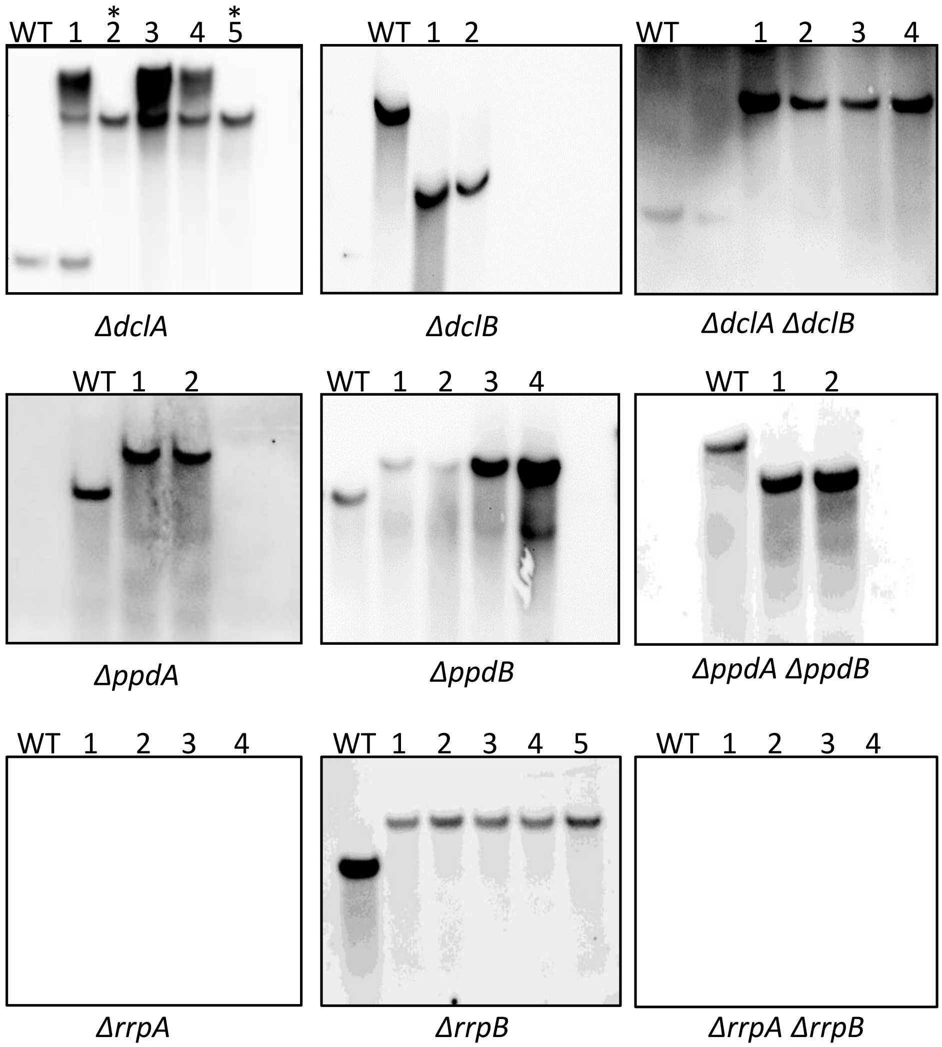

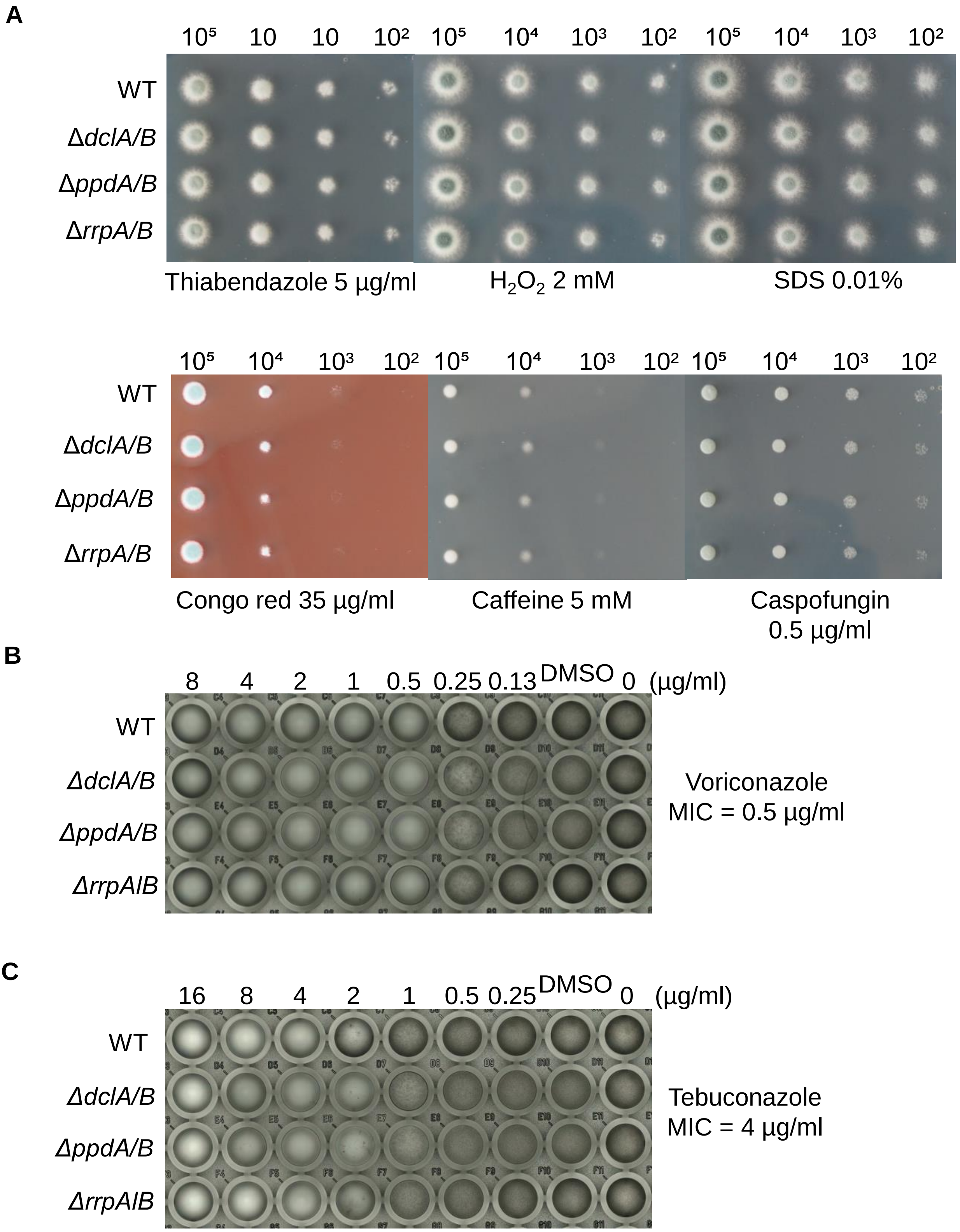

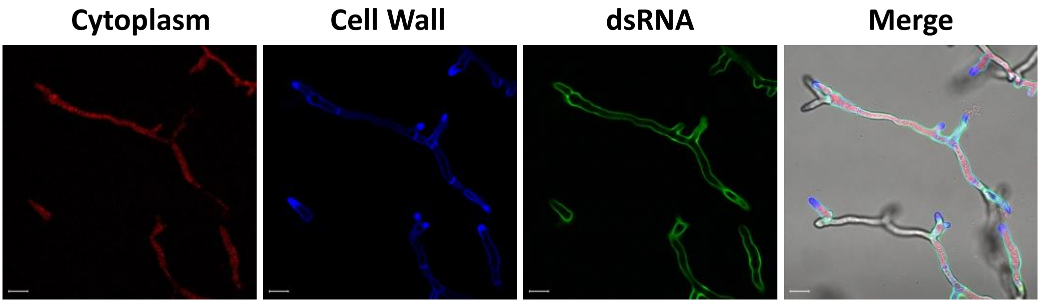

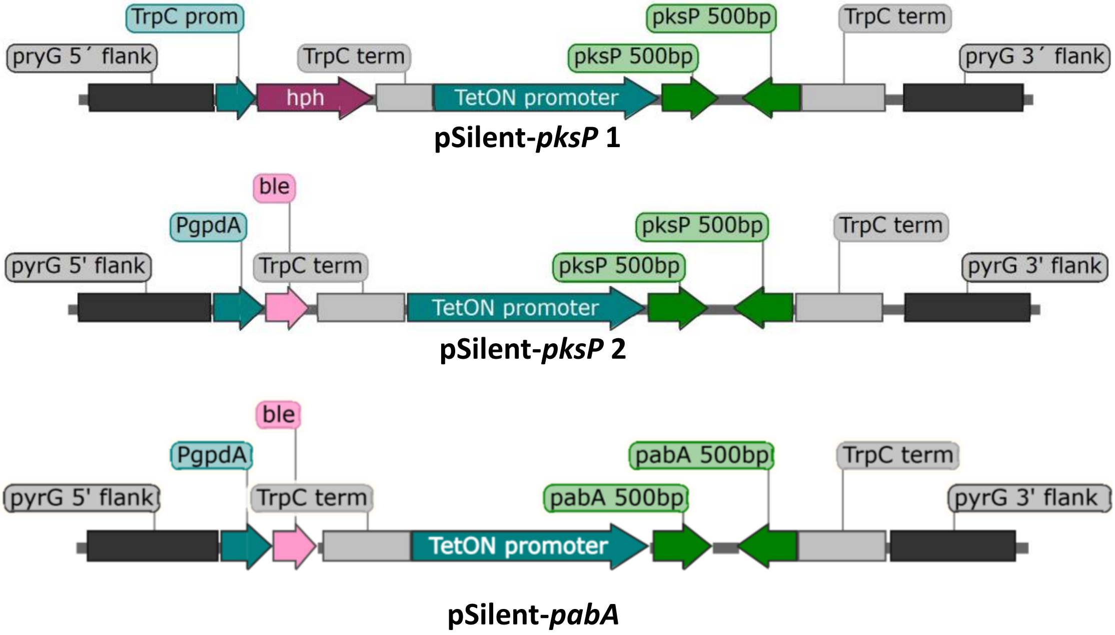

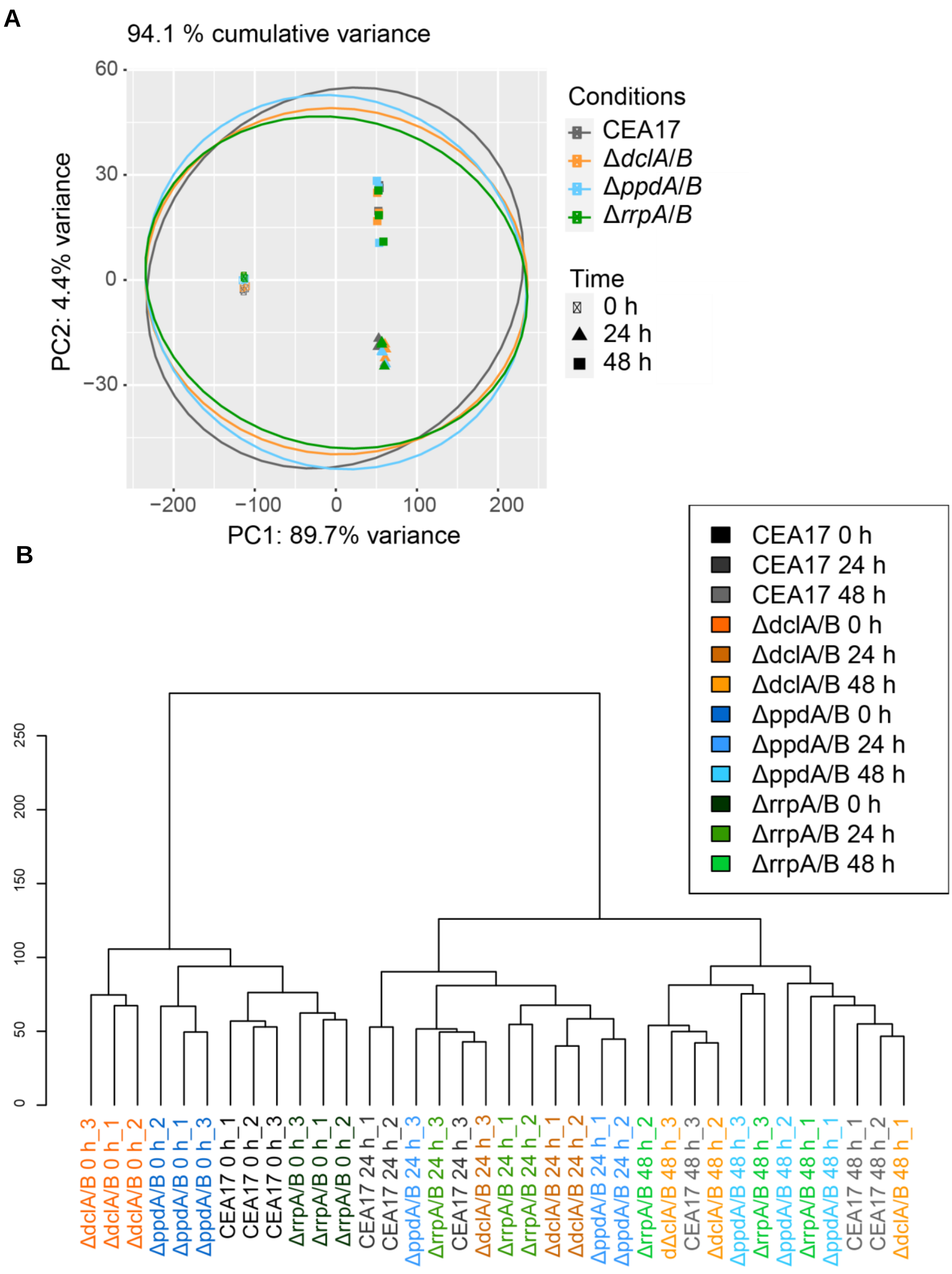

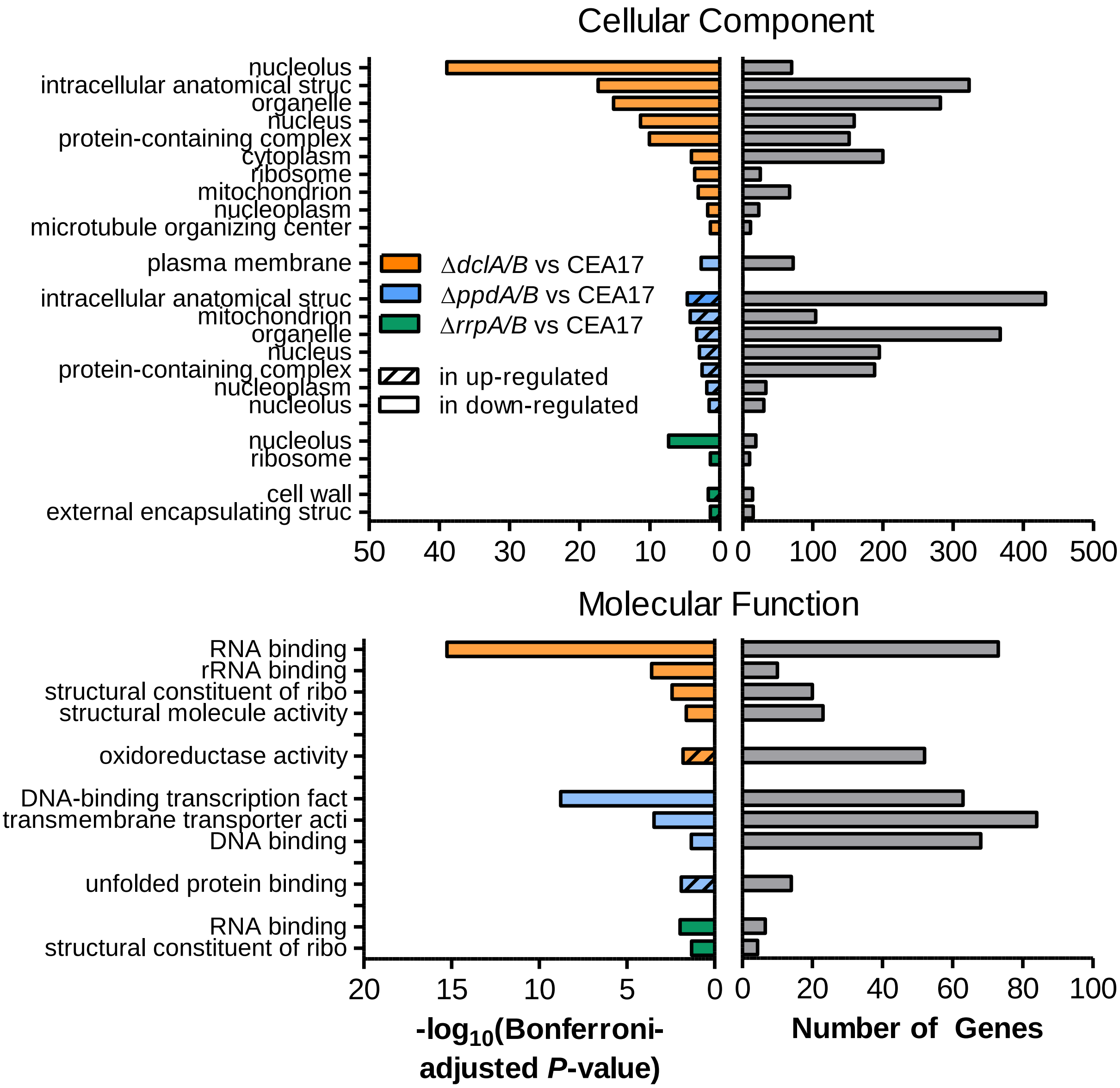

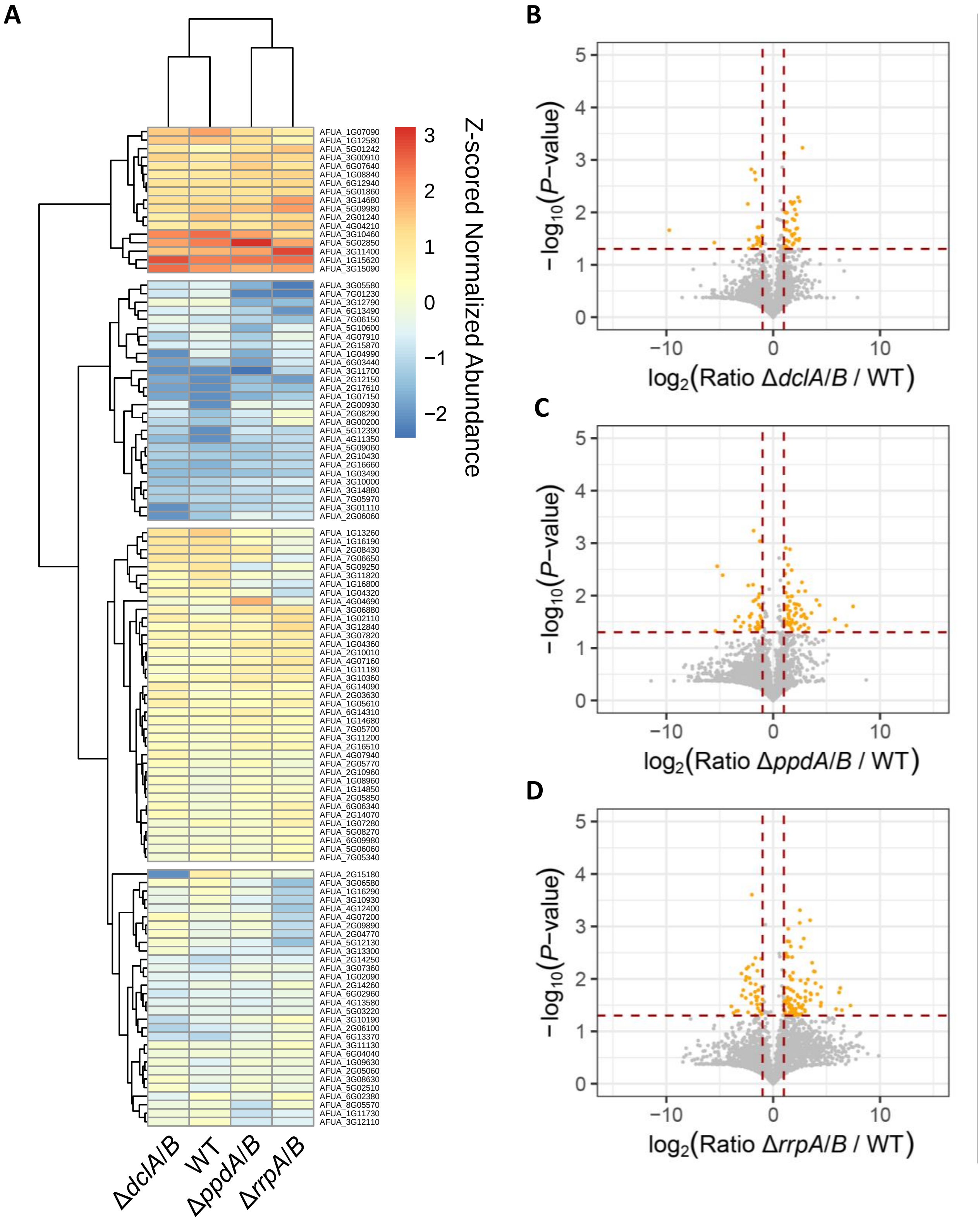

#### Figure S8

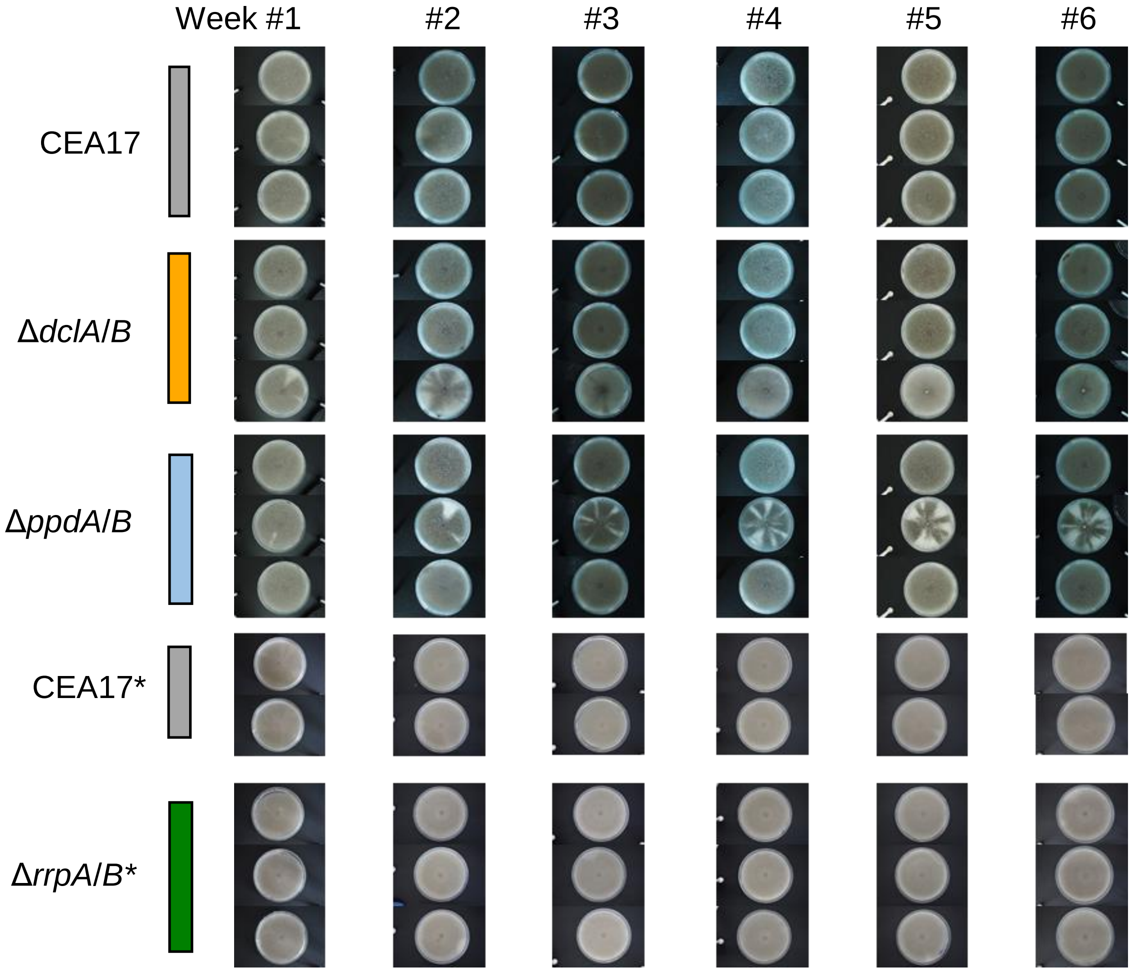
